# Functional interactomes of the Ebola virus polymerase identified by proximity proteomics in the context of viral replication

**DOI:** 10.1101/2021.07.20.453153

**Authors:** Jingru Fang, Colette Pietzsch, George Tsaprailis, Gogce Crynen, Kelvin Frank Cho, Alice Y. Ting, Alexander Bukreyev, Juan Carlos de la Torre, Erica Ollmann Saphire

**Affiliations:** La Jolla Institute for Immunology, La Jolla CA 92037; Department of Immunology and Microbiology, Scripps Research, La Jolla CA 92037; Department of Pathology, University of Texas Medical Branch, Galveston, TX 77550; Galveston National Laboratory, University of Texas Medical Branch, Galveston, TX 77550; Proteomics Core, Scripps Research, Jupiter, FL 33458; Bioinformatics and Statistics Core, Scripps Research, Jupiter, FL 33458; Cancer Biology Program, Stanford University, Stanford, CA 94305; Department of Genetics, Department of Biology and Department of Chemistry, Stanford University, Stanford, CA 94305; Chan Zuckerberg Biohub, San Francisco, CA 94158; Department of Microbiology and Immunology, University of Texas Medical Branch, Galveston, TX 77550

## Abstract

Ebola virus (EBOV) critically depends on the viral polymerase to replicate and transcribe the viral RNA genome in the cytoplasm of host cells, where cellular factors can antagonize or facilitate the virus life cycle. Here we leveraged proximity proteomics and conducted an siRNA screen to define the functional interactome of EBOV polymerase. As proof-of-principle, we validated two cellular mRNA decay factors from 35 identified host factors: eukaryotic peptide chain release factor subunit 3a (eRF3a/GSPT1) and up-frameshift protein 1 (UPF1). Our data suggest that EBOV can subvert restrictions of cellular mRNA decay and repurpose both GSPT1 and UPF1 to promote viral replication. Treating EBOV-infected human hepatocytes with a drug candidate that targets GSPT1 for degradation significantly reduced viral RNA load and particle production. Our work demonstrates the utility of proximity proteomics to capture the functional host-interactome of the EBOV polymerase and to illuminate host-dependent regulations of viral RNA synthesis.

## INTRODUCTION

Ebola virus (EBOV) is a member of the *Filoviridae* family in the order of *Mononegavirales*. Several ebolaviruses cause unpredictable sporadic outbreaks of Ebola virus disease (EVD) that are associated with high mortality. The 2013-2016 epidemic emerged in Western Africa far from sites of previously known EBOV outbreaks, entered urban populations, and caused over 11,000 deaths. EBOV re-emerged in 2018 and again in 2020 and 2021 (WHO and CDC). EVD survivors of EBOV infections can experience long-term sequelae (Adekanmbi et al., 2021; Eghrari et al., 2021; Xu et al., 2019), and may harbor virus in immune-privileged sites, posing a risk of transmitting the virus months to years after recovery (Dokubo et al., 2018; Subissi et al., 2018). The Ebola Zaire Vaccine, Live (Ervebo) and an EBOV-specific monoclonal antibody cocktail (Inmazeb), have received FDA approval for prevention and treatment, respectively, of EBOV infection. However, no small molecule antivirals, which can be delivered at lower cost, and can suppress viral replication in immune-privileged sites, are available to treat EVD. Further, small molecule antivirals may have the advantage to be broad-spectrum against multiple filoviruses if targeting a conserved viral protein.

The EBOV genome is a single-stranded, negative-sense, multicistronic RNA molecule contains seven adjacent genes separated by gene borders. The largest gene, L, encodes the catalytic subunit of EBOV RNA-dependent RNA polymerase (RdRp), which is essential for replication and expression of the viral RNA genome. L and the viral polymerase cofactor VP35 form the functional virus polymerase that acts on the EBOV genome (vRNA). In addition, EBOV encodes a viral transcription activator, VP30, which assists EBOV transcription. The vRNA coated by the viral nucleoprotein (NP) and associated with the RdRp, form the viral ribonucleoprotein complex (vRNP) (Mühlberger, 2007).

Following cell entry, EBOV delivers its vRNP into the host cell cytoplasm where viral replication and transcription occur. In the transcriptase mode, EBOV polymerase starts at a single promoter located at the 3’-end of the genome, and enters each gene at the gene start (GS) signal and discontinuously proceeding through the multicistronic viral genome, stopping at each gene end (GE) signal. Both GS and GE are conserved across all gene borders. At each GE, EBOV polymerase can either reinitiate transcription at the adjacent GS signal of the downstream gene, or dissociate and return to the same 3’ viral promoter element to initiate a new round of transcription. Due to this transcription attenuation at each gene border, EBOV polymerase synthesizes mostly monocistronic viral mRNAs that are present in a progressively decreasing amounts relative to the distance from the promoter (Hume and Muhlberger, 2019). The EBOV polymerase can also adopt a replicase mode, using vRNA as a template to produce a full-length antigenome (also complementary RNA; cRNA), which in turn serves as the template for synthesis of large amounts of progeny vRNA (Mühlberger, 2007).

The mechanisms by which EBOV executes transcriptase and replicase activities are not fully understood, partly due to a lack of structural insights on the full-length EBOV polymerase. Coordination of multiple domains with distinct enzymatic functions and conformational rearrangement of different domains can endow RdRps of negative-strand RNA viruses with different functions (Te Velthuis et al., 2021). Viral trans-factors can also facilitate function switching (Fearns and Collins, 1999; Muhlberger et al., 1999). For instance, the EBOV transcription factor VP30 can recognize non-coding, cis-regulatory sequences within the viral genome and contribute to transcriptase activity of EBOV polymerase (Biedenkopf et al., 2013; Biedenkopf et al., 2016; Ilinykh et al., 2014).

Here, we asked what role cellular factors might play in modulating distinct steps of viral RNA synthesis mediated by the EBOV polymerase. A previous study examined cellular factors that interact with the EBOV polymerase, but the results were based on over-expression of L without needed viral cofactors (Takahashi et al., 2013), and thus may not have captured the complete functional polymerase interactome.

We used recently developed proximity labeling technologies (Branon et al., 2018; Cho et al., 2020) to characterize the cellular interactomes of EBOV polymerase in the context of viral RNA synthesis in living cells. We verified the functional role of high-confidence hits identified in the interactomes using a high-content imaging-based siRNA screen with authentic EBOV infection. As a proof of concept, we further characterized two of the hits, eukaryotic peptide chain release factor subunit 3a (eRF3a/GSPT1) and the up-frameshift protein 1 (UPF1) and we uncovered their unexpected roles in EBOV infection. Our work elucidates a network of host factors that interact with EBOV polymerase and participate in the EBOV life cycle. Follow-up studies on these host factors can provide new insights into EBOV replication and illuminate novel therapeutic targets for small molecule antiviral development or drug repurposing.

## RESULTS

### Generating a functional EBOV polymerase with proximity labeling activity

We selected TurboID, an engineered promiscuous biotin ligase, optimized for biotinylation of exposed lysine residues on proteins within a ∼10 nm radius (Branon et al., 2018). Since EBOV polymerase (EBOV_pol) requires both L and VP35 proteins to function, we used the split-TurboID system, consisting of two inactive fragments, sN- and sC-TurboID (Cho et al., 2020). When brought together via protein-protein interactions, sN- and sC-TurboID reconstitute full-length TurboID and its biotin ligase activity.

To construct EBOV L-sNTurboID, we inserted sNTurboID into a site of EBOV L known to tolerate insertion of mCherry (Hoenen et al., 2012). For EBOV VP35-HA-sCTurboID, we fused the sCTurboID and a HA tag to the EBOV VP35 C terminus, thus preserving the N terminus and internal oligomerization domain, which are implicated in binding to NP and L, respectively, and which are both needed to support viral RNA synthesis (Leung et al., 2015; Moller et al., 2005). Interaction of EBOV L with VP35 allows trans-complementation of the associated split-TurboID fragments and biotinylation of neighboring proteins. In addition, the presence of the EBOV transcription factor VP30 promotes conversion of EBOV_pol from a replicase to a transcriptase (Biedenkopf et al., 2013), and may modulate the polymerase interactome (**Figure 1A**).

**Figure 1.**
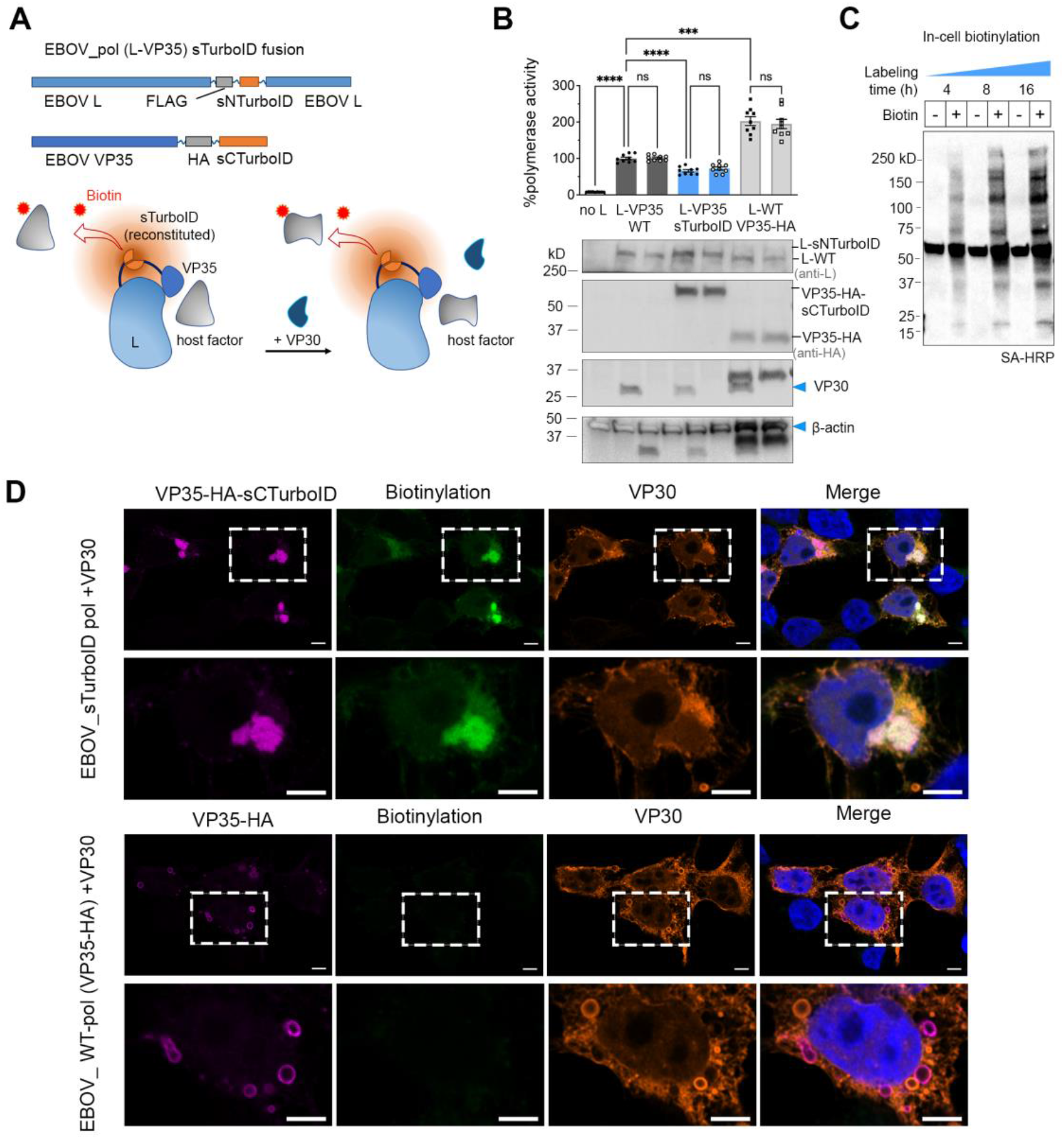
Generating a functional EBOV polymerase with proximity-labeling activity. **(A)** Proximity biotinylation by the EBOV polymerase (EBOV_pol) split-TurboID fusion with and without VP30. **(B)** Activity and expression of EBOV_pol split-TurboID fusion (L-VP35 sTurboID) compared to controls (L-VP35 WT and L-WT VP35-HA) measured in EBOV minigenome (MG) assay. % Polymerase activity is determined by normalizing luciferase activity in each sample to that of L-VP35 WT control. Background of MG activity is represented by the “no L” control. Results from three independent experiments with triplicate wells were analyzed by Welch’s ANOVA with Dunnett’s multiple comparisons test comparing each condition with controls (ns, not significant; *, *p < 0*.*05*; **, *p < 0*.*01*; ***, *p < 0*.*001*). Values are means ± SEM. Representative western blot showing expression of EBOV L-variants, VP35-variants, VP30 and β-actin in HEK 293T lysates. The blot was sliced into two parts to sequentially probe for different target proteins. **(C)** Streptavitin (SA)-blot showing biotin ligase activity of L-VP35 sTurboID in the context of the EBOV MG system. **(D)** Confocal immunofluorescence images of L-VP35 sTurboID mediated proximity biotinylation in cells co-expressing the EBOV MG system. VP35-HA and VP35-HA-sNTurboID were all detected by anti-HA. EBOV VP30 was detected by anti-VP30. Biotinylated proteins were detected by streptavidin-AF488. Scale bar: 5 μm. Representative images from two independent experiments. Zoomed areas are outlined with a white rectangle in the original image. See also Figure S1.

We used an established EBOV viral minigenome (MG) system (Jasenosky et al., 2010a) (**Figure S1**) to confirm that EBOV_pol split-TurboID fusion is functionally active. This MG system recapitulates viral RNA synthesis using activity of a Firefly luciferase reporter as a comprehensive measure of MG replication, transcription, and translation of the MG reporter transcript. As a control, we used unmodified wild-type VP35 (VP35-WT) and wild-type L (L-WT). We measured polymerase activity in the presence or absence of VP30, using the activity of WT EBOV_pol (L-VP35 WT) in the presence of VP30 for normalization. We additionally included another control with L-WT and C-terminal HA-tagged VP35 (VP35-HA), which can be compared with VP35-HA-sCTurboID for VP35 expression. The EBOV_pol split-TurboID fusion (L-VP35-sTurboID) retains 65% of WT polymerase activity (L-VP35-WT); 35% of L-WT and VP35-HA control, without noticeable difference in expression levels of L proteins. VP35-sCTurboID was expressed at higher levels than the VP35-HA control. Surprisingly, withdrawal of VP30 did not affect MG activity using either WT or split-TurboID tagged EBOV_pol (**Figure 1B**). This suggests that, in contrast to other EBOV MG systems (Biedenkopf et al., 2016), the EBOV MG construct we used here permits VP30-independent transcription, a non-canonical transcription mode (Weik et al., 2002). This feature of the EBOV MG being used allowed us to probe a context-dependent viral polymerase interactome without VP30, and downstream validation of VP30-dependent roles of interactors of interest.

We validated the proximity labeling activity by detecting biotinylated proteins in cells expressing EBOV_pol split-TurboID fusion together with EBOV MG system components. EBOV_pol split-TurboID fusion produced, in a time-dependent manner, a broad range of biotinylated proteins in the presence of exogenous biotin (**Figure 1C**).

We used confocal microscopy to detect biotinylation by EBOV_pol split-TurboID fusion in fixed cell specimens. HEK 293T cells transfected with the EBOV MG system, either with the split-TurboID fusion or the control EBOV_pol (with L-WT and VP35-HA) were labeled with biotin for one hour at one day post-transfection. Because the lack of an EBOV L-specific monoclonal antibody, we used the polymerase cofactor VP35 localization as a proxy for EBOV_pol localization. VP35-sCTurboID fluorescence signals were confined to viral inclusion bodies and overlapped with biotinylation signals. The VP35-HA control was localized only to the edges of inclusion bodies and no endogenous biotinylation signal was observed (**Figure 1D**). These different localization patterns of VP35 may be due to the higher expression level of VP35-sTurboID relative to VP35-HA or partitioning of other viral proteins in the EBOV inclusion body, as both phenotypes (i.e., inclusion-filling and inclusion-edge) occur in EBOV-infected cells (Nanbo et al., 2013). Most EBOV VP30 did not co-localize with VP35-positive inclusion bodies, but instead formed its own inclusions, a finding consistent with the VP30-independent activity of the EBOV MG system used.

### Proximity proteomics to define EBOV polymerase interactomes

We transfected HEK 293T cells with the EBOV MG system components, including wild-type EBOV_pol (WT_pol) or EBOV_pol split-TurboID (sTurboID_pol), followed by biotin labeling. Transfected cells were lysed, and biotinylated proteins were captured with streptavidin (SA) beads that were then subjected to on-bead trypsin digestion. Digested peptides were labeled with unique tandem-mass tags (TMTs) for quantitative proteomics (**Figure 2A**).

**Figure 2.**
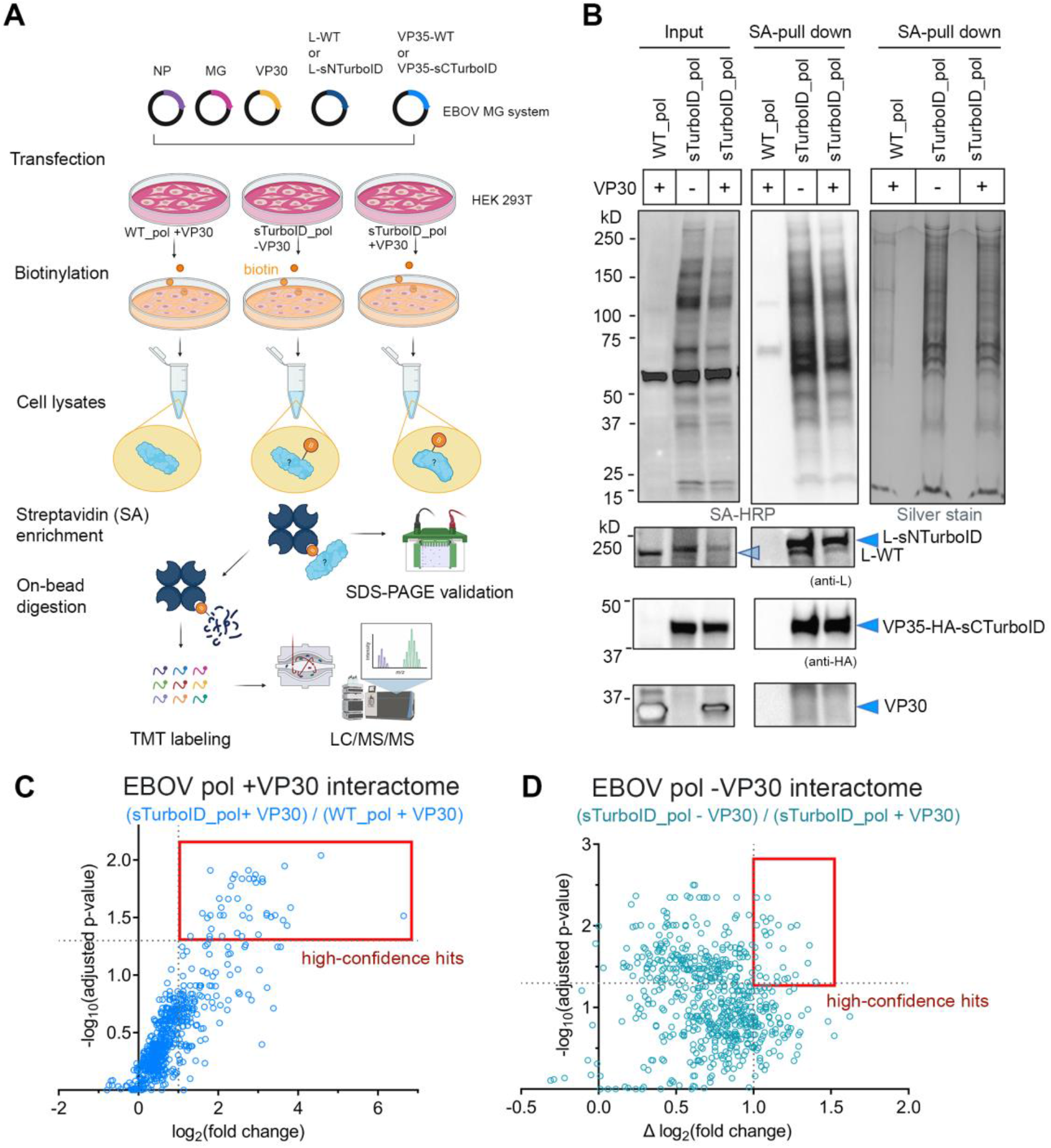
Proximity proteomics to define the EBOV polymerase interactomes. **(A)** Proximity proteomics workflow. **(B)** Validation of streptavidin (SA) enrichment of EBOV_pol split-TurboID (sTurboID_pol) mediated proximity biotinylation. “Input”: total lysates; “SA-pull down”: biotinylated proteins enriched by SA-beads. Representative blots and gels from three biological replicates are shown. All samples were split into three parts that were loaded onto individual gels used for streptavidin-HRP blot, to probe EBOV proteins, and for silver staining. The blot used to probe EBOV proteins was sliced into three parts that were imaged separately. Scatter plots showing interactors of EBOV polymerase + VP30 **(C)** and EBOV polymerase -VP30 **(D)** in HEK 293T cells. The degree of enrichment for individual proteins is quantified by fold-change and statistical confidence of enrichment (log-transformed, adjusted P value). See also Table S1.

To assess enrichment efficiency, biotinylated proteins bound to SA beads (SA-pull down) from equal amounts of starting material (Input) was eluted in SDS loading buffer and analyzed by western blot (**Figure 2B**). We observed enrichment of biotinylated proteins specific to samples with sTurboID_pol expression compared to those with WT_pol expression. The addition of VP30 to sTurboID_pol slightly suppressed expression of both sTurboID_pol and VP30 itself, resulting in fewer biotinylated proteins. Both components of EBOV sTurboID_pol were detected in the SA-pull down, suggesting that self-biotinylation occurred. Unexpectedly, the SA-pull down contained no VP30, indicating that it may not directly associate with EBOV L or VP35, or that its spatial localization was outside the labeling radius of sTurboID_pol. Consistent with this finding, we saw no appreciable colocalization of VP30 with VP35-positive inclusion bodies (**Figure 1D**).

To analyze EBOV_pol interactomes, we assessed host proteins identified in the proximity interactome by two criteria: degree of enrichment compared to the control proteome (WT_pol), and the corresponding statistical confidence. We normalized the abundance ratio for each protein in the sTurboID_pol sample to that in the WT_pol sample and determined the value of fold-change. We then performed multiple comparisons across three biological replicates and determined adjusted *p*-values (adjusted to False Discovery Rate of 0.05) to identify proteins enriched in the sTurboID_pol interactome. High-confidence hits were selected using a threshold of adjusted *p-value* < 0.05 and log_2_(fold change) > 1. We identified 43 hits that interact with the EBOV polymerase in the presence of VP30 (**Figure 2C**). Another 28 hits were found in the absence of VP30 (≥ 2-fold higher abundance ratio in the sTurboID_pol -VP30 vs the sTurboID_pol +VP30 samples) (**Figure 2D**).

### Effect of siRNA targeting high-confidence EBOV polymerase interactors on viral infection

Next, we used an siRNA-based functional screen (Fang et al., 2018) to assess how the EBOV polymerase-interacting protein candidates affected EBOV infection. We transfected Huh7 cells with individual siRNAs targeting each of 64 high-confidence hits and subsequently infected them with recombinant EBOV-eGFP (Towner et al., 2005). We determined the percentage of GFP-positive cells, as an indicator of EBOV infection, at 48 hours post-infection (hpi) (**Figure 3A**). We excluded targets for which siRNA transfection altered cell count by one standard-deviation from the average cell count, and considered true hits as those having at least two independent siRNA-KD resulting in enhanced or decreased viral infection across two biological replicates.

**Figure 3.**
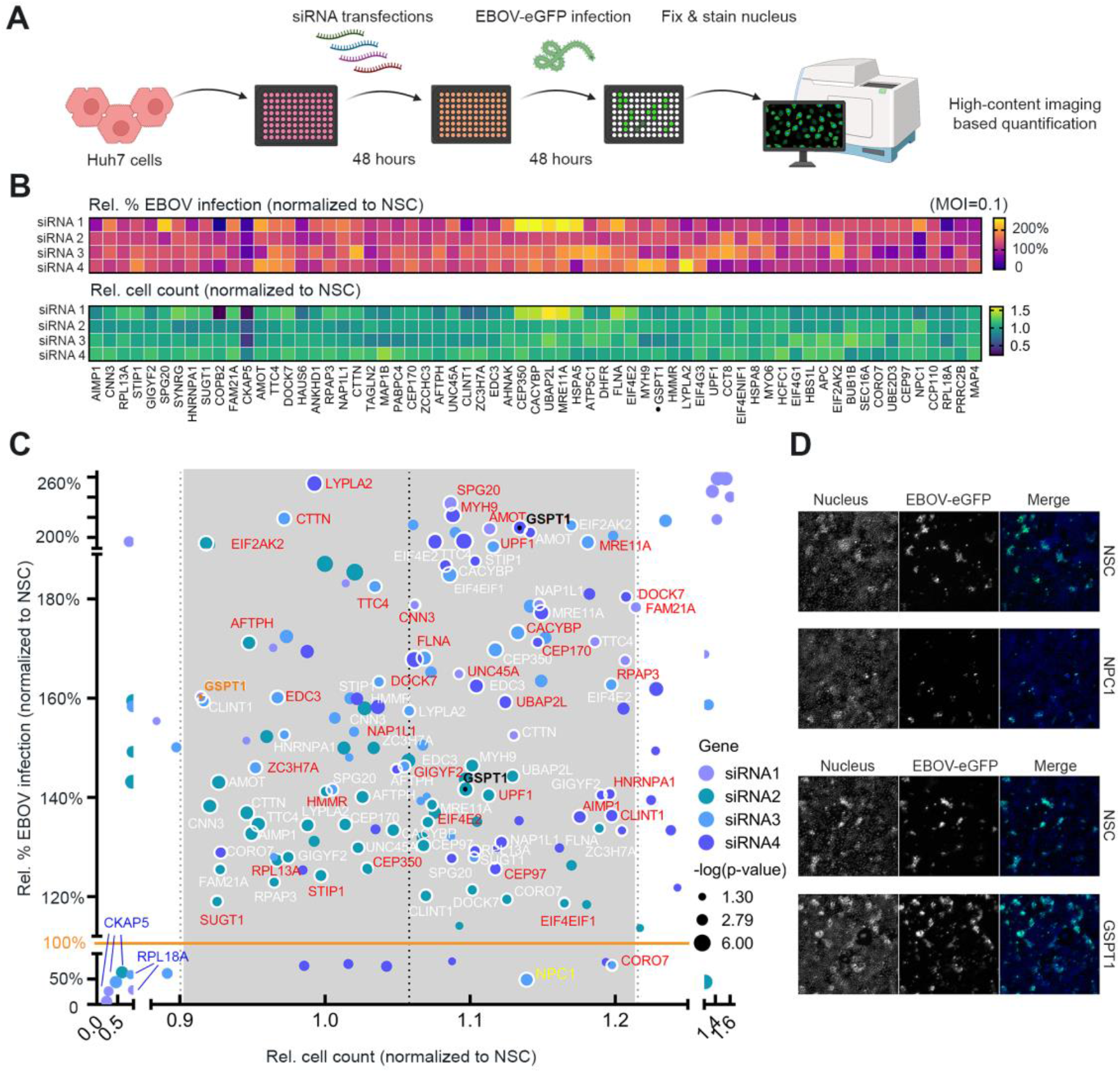
Effect of siRNA targeting high-confidence EBOV polymerase interactors on viral infection. **(A)** siRNA screen with authentic EBOV infection. **(B)** Normalized values displayed in heat maps of relative percentage of EBOV infection and relative cell count. Each value is the mean of triplicate wells transfected with the same siRNA. Multiple unpaired t-tests were performed to determine the statistical significance of siRNA-mediated changes on % EBOV infection compared to that of NSC. The determined P values were log-transformed and those having *p* > 0.05 were excluded. **(C)**. Individual genes for which siRNAs significantly affected infection are shown as data points in a bubble plot; data point colors and sizes correspond to the siRNA tested and log-transformed P value, respectively. Data points with multiple siRNAs significantly modulating EBOV infection are considered as true hits and are outlined in white with the corresponding gene also labeled in white. One label of each true hit is shown in red for display purposes. NPC1 used as a positive control is labeled in yellow. GSPT1 is labeled in black. **(D)** Representative images of selected Huh7 monolayers at 48 hours post-infection (hpi). One representative result from two rounds of siRNA screens using different multiplicities of infection/MOI (PFU/cell) is shown. See also Figure S2-S3.

Most hits appeared to be antiviral at 48 hpi, shown by an increased percentage of EBOV infection in the presence of siRNA-KD (**Figure 3B** and **S2A**). Limited (10-30% of cells) infection rates for cells transfected with non-silencing control (NSC) siRNA may have favored detection of enhanced infection over reduced infection. Our results are supported by stringent statistical metrics (two biological replicates with different multiplicities of infection/MOIs, four independent siRNAs per target) and guided by quantitative parameters of the resulting phenotype (i.e., Relative % infection and cell count) (**Figure S2B**). We identified 35 potential EBOV-specific antiviral factors (**Figure 3C** and **3D, S4**). Knockdown of CKAP5 or RPL18A reduced both EBOV infection and cell count, which prevented their consideration as pro-viral factors. In contrast, knockdown of the EBOV entry receptor NPC1 (positive control) inhibited EBOV infection by nearly 50%, without significantly affecting the cell count (**Figure 3D**, top panel).

Based on these results, we generated an EBOV polymerase interactome network by clustering all functional polymerase interactors according to their STRING-classified biological processes (Szklarczyk et al., 2019). Around 80% of the interactors have not been previously described, many of them are involved in critical cellular pathways relevant to the EBOV life cycle (**Figure 4A**). Twelve interactors were reported in other EBOV-protein interactomes (**Figure 4B**) (Fan et al., 2020; Garcia-Dorival et al., 2016; Morwitzer et al., 2019; Spurgers et al., 2010; Takahashi et al., 2013).

**Figure 4.**
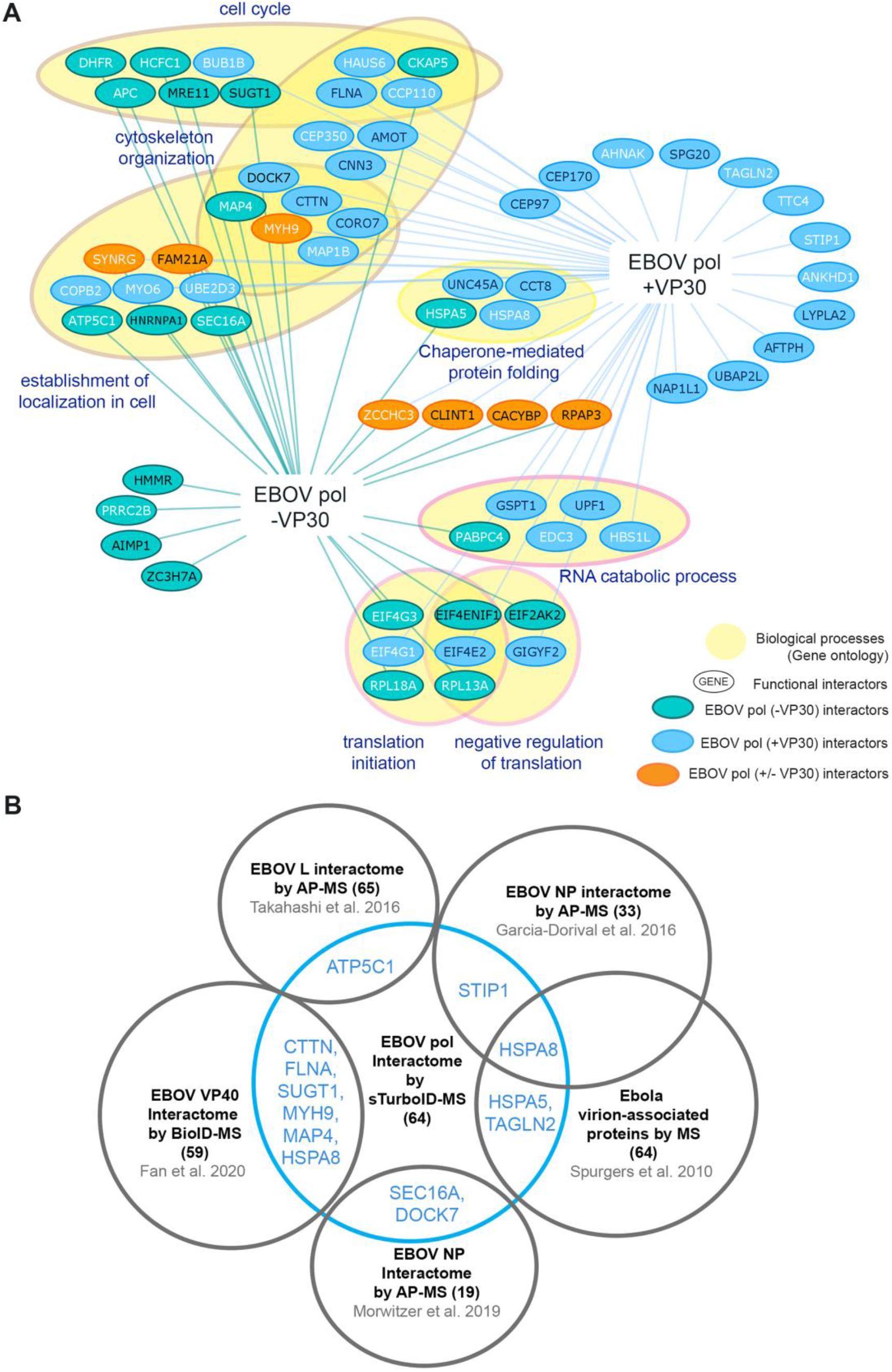
Proximity interactomes of EBOV polymerase in the presence and absence of EBOV VP30. **(A)** EBOV polymerase interactomes identified by proximity proteomics. Node (colored ovals): high-confidence proteomic hit. Edge (colored lines): identified protein-protein interaction. Nodes having two edges are shared by multiple interactomes. Black text inside nodes denotes genes that are functionally validated by siRNA screens. Selected nodes are clustered based on enriched biological processes using STRING. **(B)** A Venn diagram comparing EBOV polymerase interactors with reported interactors of EBOV proteins or virions.

### GSPT1 is functionally relevant to EBOV RNA synthesis

Two functional polymerase interactors, eukaryotic peptide chain release factor subunit 3a (eRF3a/GSPT1) and up-frameshift protein 1 (UPF1), are key components of the cellular nonsense mRNA decay (NMD) pathway. UPF1 restricts infection by multiple RNA viruses (May and Simon, 2021), whereas GSPT1, also a translation termination factor (Zhouravleva et al., 1995), has not been implicated in viral infections. Thus, as a proof-of-principle, we examined the role GSPT1 on EBOV infection.

Using co-immunoprecipitation (co-IP) assays of HEK 293T cells co-expressing an N-terminal Flag-tagged GSPT1 (long isoform, 68.7KDa) and EBOV_pol components, either individually or in combination, we found that in the presence of VP35-WT, L-WT consistently co-IPs with Flag-GSPT1 (**Figure 5A)**. VP30 was present in the same co-IP protein complex when co-expressed with L-VP35. We observed a pronounced degradation product of L-WT in the input and unbound fraction but not the co-IP, suggesting that the Flag-GSPT1 and EBOV L-WT interaction requires full-length L (or at least those regions otherwise subjected to degradation). In contrast, Flag-GSPT1 did not co-IP with VP35 (**Figure 5B**) or VP30 (**Figure 5C**) in the absence of L, indicating that GSPT1 interacts with EBOV_pol via interacting with L protein. GSPT1 is not known to have RNA-binding activity. Therefore, the GSPT1-EBOV_pol interactions are likely not due to indirect interactions bridged by RNA molecules.

**Figure 5.**
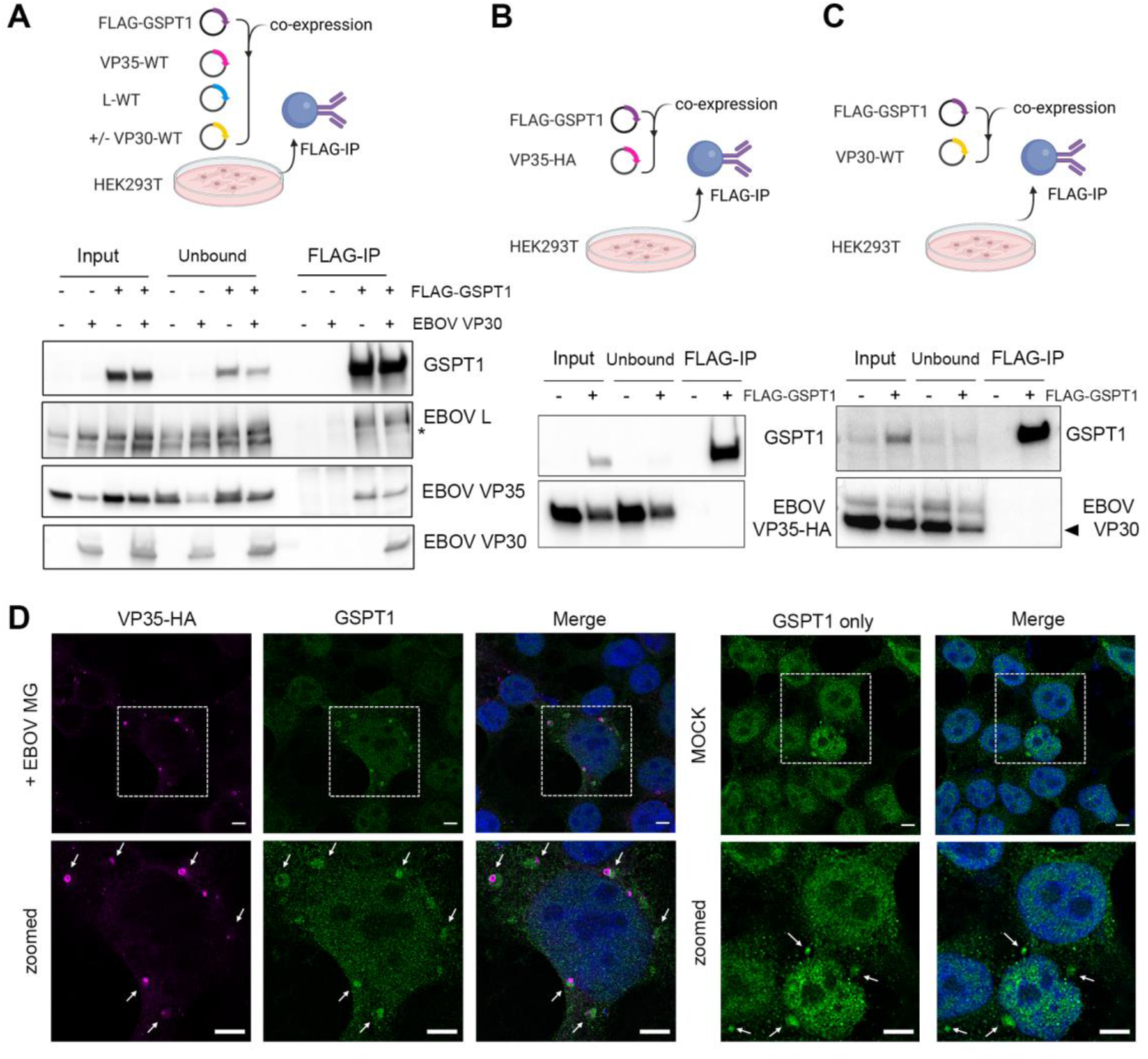
GSPT1 associates with Ebola virus polymerase. Anti-Flag pull-down of GSPT1 complexes. **(A)** Expression of EBOV polymerase complex (L-VP35) +/-VP30. **(B)** VP35 in the absence of L. **(C)** VP30 in the absence of L. Representative results from two experiments are shown. Each blot was sliced into several parts to probe for different target proteins and imaged separately. **(D)** Confocal immunofluorescent analysis of GSPT1 and EBOV_pol (by VP35-HA) localization in the context of the EBOV MG system in HEK 293T cells. Arrowhead: sites where endogenous GSPT1 localizes next to viral proteins. The bottom panel shows the subcellular distribution of GSPT1 alone as a negative control. Representative images from two experiments are shown. See also Figure S5.

We next examined whether endogenous GSPT1 interacts with EBOV_pol in a cellular context that recapitulates viral RNA synthesis. In HEK 293T cells transfected with EBOV MG system, we observed a specific pattern of endogenous GSPT1 clusters in the cytoplasm that appeared to contact EBOV VP35-positive inclusion bodies (**Figure 5D**, left panel). In control cells, endogenous GSPT1 had a mostly diffuse nucleocytoplasmic distribution (**Figure 5D**, right panel). We therefore examined the functional consequences of GSPT1 association with EBOV_pol using the EBOV MG assay. Consistent with the siRNA screen results, over-expression or GSPT1 knockdown (KD) significantly decreased and increased, respectively, EBOV MG activity. Meanwhile, changes in GSPT1 expression did not affect expression of a control eGFP-reporter (**Figure S3A and S3B**). Together, our data suggest that GSPT1 is a *bona fide* EBOV polymerase interactor that may participate in EBOV viral RNA synthesis.

### EBOV hijacks GSPT1 to facilitate transcription termination

To further characterize the phenotype of GSPT1-KD in the context of EBOV infection, we next followed the effect of GSPT1-KD on multi-step EBOV growth kinetics in Huh7 cells. GSPT1-KD persisted for seven days post-siRNA transfection, supporting that any reduction in endogenous GSPT1 expression would be sustained throughout the four-day viral growth curve experiments (**Figure S4**). We infected GSPT1-KD cells with EBOV and measured viral titers in the cell supernatant on four consecutive days.

GSPT1-KD significantly changed EBOV growth kinetics in Huh7 cells during the four-day time course (**Figure 6A**). On day 1, GSPT1-KD mildly increased EBOV titers by two-fold relative to the non-silencing control (NSC). However, from 2-to 4-day post-infection (dpi), GSPT1-KD led to reduced EBOV titers. At 4 dpi, we observed a significant decrease in EBOV vRNA accumulation (**Figure 6B**), fewer viral infected cells (**Figure 6C**), and reduced levels of viral proteins in cells with GSPT1-KD (**Figure 6D)**. However, we also found universally increased levels of EBOV mRNA accumulation upon GSPT1-KD at 4 dpi (**Figure 6E**), which did not translate into any increased viral protein accumulation. We hypothesized that the concurrent increase in EBOV mRNAs and decrease in EBOV protein accumulation associated with GSPT1-KD could reflect aberrant viral transcription involving readthrough mRNA synthesis. This type of aberrant transcription occurs when the viral polymerase reads-through the transcription termination signal of an upstream gene to continue transcribing the downstream gene unit (Brauburger et al., 2014). Products of readthrough transcription are polycistronic mRNAs that contain the sequence of gene borders, rendering the mRNAs incompetent for cap-dependent translation. Enhanced readthrough transcription could thus account for an apparent increase in EBOV mRNA levels, that is associated with fewer transcription attenuation steps, accompanied by decreased translation of the downstream viral genes in the readthrough mRNAs.

**Figure 6.**
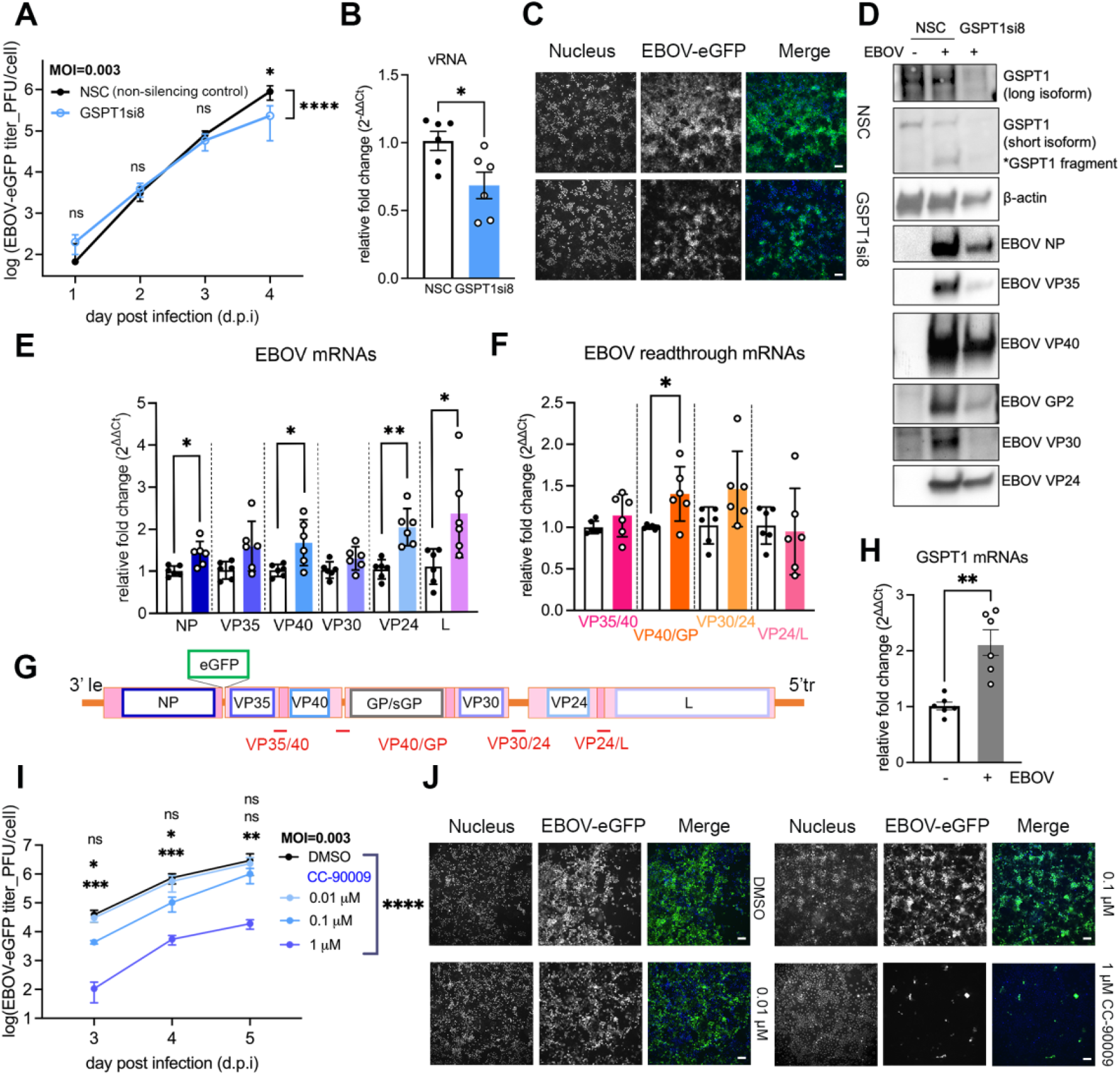
EBOV subverts GSPT1 restriction and hijacks GSPT1 to facilitate termination of viral transcription. **(A)** Effect of GSPT1-KD on EBOV viral growth kinetics in Huh7 cells (MOI=0.003 PFU/cell). Results of two independent experiments with triplicates are plotted as mean ± SD. Two-way ANOVA analysis of log-transformed viral titers was performed to determine the statistical significance of the effect of GSPT1 depletion by siRNA on viral growth kinetics. Multiple comparisons were performed at each time point between non-silencing control (NSC) and GSPT1si8. **(B)** Effect of GSPT1-KD on EBOV vRNA (genome) accumulation at 4 days post-infection (dpi). **(C)** Representative images of EBOV-infected Huh7 monolayer with and without GSPT1-KD at 4 dpi. Scale bar: 100 µm. **(D)** Western blot analysis of GSPT1 protein, loading control (β-actin), and viral protein levels in lysates from EBOV-infected Huh7 cells harvested at 4 dpi. Representative blots from one experiment are shown. Equal volumes of inactivated lysate were pooled from triplicates with the same treatment. Due to the low expression level of L protein, we could not consistently detect full-length L in inactivated lysate. **(E)** Effect of GSPT1-KD on the level of **(F)** individual EBOV mRNA and **(G)** EBOV readthrough mRNAs. Locations of each readthrough sequence on the viral genome are marked as red bars in (G). **(G)** Schematic of the EBOV-eGFP genome used in the viral growth kinetics experiment. Orange bars: non-coding terminal and intergenic regions. Pink box: gene unit with the coding sequence outlined in a different color. For all bar graphs, results from two independent experiments with triplicates were analyzed by using Welch’s t test under each condition compared to controls (*, *p < 0*.*05*; **, *p< 0*.*01)*. Values are means ± SD. White bar: NSC; colored bar: GSPT1si8. **(H)** Effect of EBOV infection on endogenous GSPT1 expression in Huh7 cells, compared to no infection. **(I)** Effect of small molecule CC-90009 on EBOV viral growth kinetics in Huh7 cells (MOI=0.003 PFU/cell). CC-90009 added 1 h after infection. Results of one experiment with triplicates are plotted as mean ± SD. Two-way ANOVA analysis of log-transformed viral titers was performed. Multiple comparisons were used to compare at each time point between DMSO and CC-90009 treatment. **(J)** Representative images of EBOV-infected, mock or drug-treated Huh7 monolayer at 5 dpi are shown. Scale bar: 100 µm. See also Figure S6-8.

Consistent with this hypothesis, the magnitude of the increase in mRNA was greater for genes located on the promoter-distal, 5’ end of the EBOV genome, compared to genes on the promoter-proximal, 3’ end at the polymerase entry site (**Figure 6F**). Accordingly, in GSPT1-KD cells, we detected higher incidence of EBOV transcription read-through in 3/4 gene borders we examined (**Figure 6G**). These results confirmed our hypothesis of enhanced transcription readthrough upon GSPT1-KD. With fewer viral genome being synthesized and fewer viral proteins being translated, our results suggest a less efficient EBOV infection in GSPT1-KD cells and in turn reduced production of infectious particles measured as EBOV titer.

### Pharmacological targeting of GSPT1 protein strongly inhibits EBOV infection

We observed that EBOV infection itself induced significant upregulation of GSPT1 transcripts (**Figure 6H**). This may partially explain the less robust siRNA-mediated GSPT1 depletion with EBOV infection (**Figure 6D**) than without EBOV infection (**Figure S4**). To overcome this hurdle, we used a small molecule degrader/PROTAC, CC-90009, known to trigger the proteasomal degradation of GSPT1 protein through tethering GSPT1 with a cellular ubiquitin ligase (Hansen et al., 2021; Surka et al., 2021). We infected (MOI = 0.003 PFU/cell) Huh7 with EBOV, and treated them with CC-90009 (**Figure S6E**). CC-90009 treatment at 1 μM potently reduced the EBOV titer in the supernatant (**Figure 6I**). We detected a dose-dependent effect of CC-90009 on levels of EBOV vRNA and mRNA (**Figure S6A and S6B**). Further, we observed reductions of GSPT1 and EBOV proteins in cells treated with CC-90009 at 1 μM, but not at 0.1 or 0.01 μM (**Figure S6D**).

The consistent appearance of a lower-molecular weight GSPT1 species (∼40kDa) that could correspond to a cleavage product of GSPT1, in both GSPT1-KD cells by siRNA and CC-90009-treated cells is interesting. This phenotype was specific to EBOV infection, highlighting a complex interplay between GSPT1 and EBOV. We also observed that CC-90009 caused a dose-dependent increase in the level of GSPT1 transcripts both with and without EBOV infection (**Figure S6C and S6F**). The underlying mechanisms for this observation remain to be determined.

In summary, our data suggest at early time points, GSPT1 may restrict EBOV infection, whereas later in viral infection, GSPT1 can be hijacked by the EBOV polymerase to facilitate critical viral RNA synthesis processes.

### EBOV can subvert the restriction by UPF1

Given that EBOV infection is not restricted by GSPT1, we asked whether this is true for another nonsense-mediated decay (NMD) factor identified in our screen, UPF1. We again evaluated the EBOV growth kinetics in Huh7 cells with siRNA-mediated UPF1-KD (**Figure S6**).

With UPF1-KD, we observed that the one-step EBOV growth curve (MOI=3 PFU/cell) was significantly altered with increasing time. EBOV titers were slightly attenuated at 18 hpi, but the viral growth recovered after 24 hpi and subsequently increased (**Figure 7A**). We also observed the multi-step EBOV growth kinetics (MOI=0.003 PFU/cell) was enhanced by two-fold in UPF1-KD cells at later time points, similar to that of GSPT1-KD (**Figure 7B**, compared to **Figure 6A**). We found a global reduction at the level of EBOV vRNA and mRNAs upon UPF1-KD at 4 dpi (MOI=0.003 PFU/cell) (**Figure 7C** and **7D**), indicating a different mechanism compared to GSPT1-KD, in which case EBOV vRNA and mRNAs were differentially affected. Consistent with the decreased level of EBOV RNAs, fewer cells were infected (**Figure 7E**) and fewer viral proteins were accumulated when UPF1 is depleted (**Figure 7F**). Unlike what we observed for GSPT1 transcripts, the level of UPF1 transcripts did not change upon EBOV infection in Huh7 cells (**Figure 7G**, compared to **Figure 6H**). These results suggest that UPF1 can only modestly limit EBOV infection and UPF1-restriction is subverted by EBOV as infection progresses.

**Figure 7.**
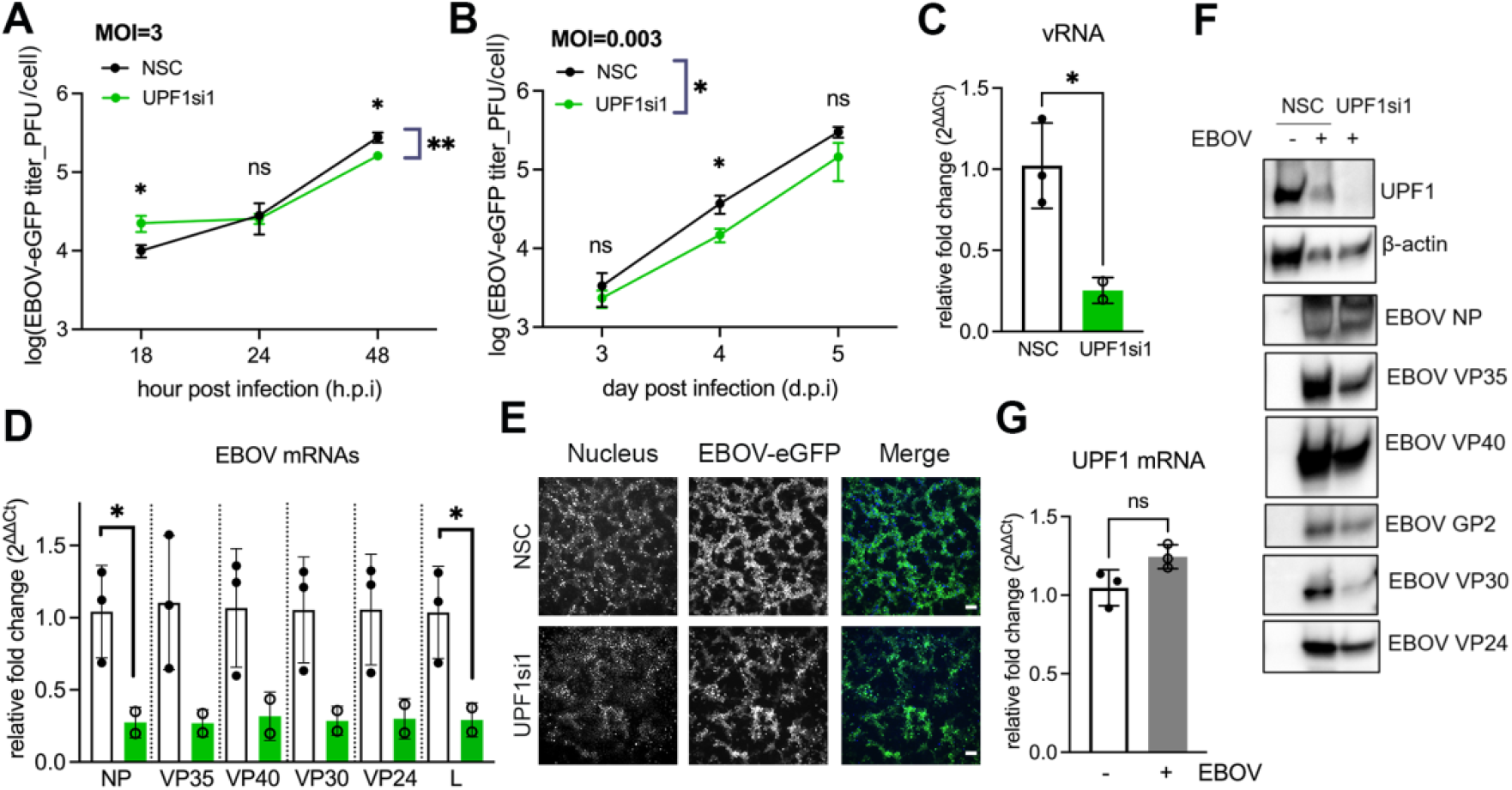
EBOV subverts UPF1-restriction and utilizes UPF1 as a pro-viral factor. Effect of UPF1-KD on EBOV viral growth kinetics in Huh7 cells at MOI=3 (PFU/cell) in early time points **(A)**; or at MOI=0.003 (PFU/cell) in later time points **(B)**. Results of one independent experiment with triplicates are plotted as mean ± SD. Two-way ANOVA analysis of log-transformed viral titers was performed to determine the statistical significance of the effect of UPF1 knockdown by siRNA on viral growth kinetics. Multiple comparisons were performed at each time point between NSC and UPF1-KD. Effect of UPF1-KD on **(C)** EBOV vRNA and **(D)** mRNAs accumulation at 4 days post-infection (dpi), MOI=0.003. For all bar graphs, results from one experiment with triplicate were analyzed by using Welch’s t test under each condition compared to controls (*, *p < 0*.*05)*. Values are means ± SD. White bar: NSC; colored bar: UPF1si1. Results for UPF1si1 includes duplicated wells. **(E)** Representative images of EBOV-infected Huh7 monolayers (MOI=0.003) at 5 dpi are shown. Scale bar: 100 µm. **(F)** Western blot analysis of UPF1, loading control (β-actin), and viral protein levels in lysates from EBOV-infected Huh7 cells (MOI=0.003) harvested at 5 dpi. Equal volumes of inactivated lysate were pooled from triplicates with the same treatment. Low expression level of L protein precluded consistent detection of the full-length L in inactivated lysate. **(G)** Effect of EBOV infection (MOI=0.003) on endogenous UPF1 transcripts in Huh7 cells, compared to no infection. Representative images of EBOV-infected Huh7 monolayers at 4 dpi are shown. See also Figure S9.

## DISCUSSION

Here we present the first systematic analysis of the EBOV polymerase-cellular interactome in the context of viral RNA synthesis. We used proximity labeling to capture the EBOV polymerase interactomes *in situ*, and performed an siRNA screen in EBOV-infected cells to identify functionally relevant EBOV polymerase interactors. We identified 64 high-confidence interactors, 31 of which were validated by our siRNA screen as functional hits. Most identified hits were antiviral, suggesting that the viral polymerase, viral RNA products, or both, are targets of host defense mechanisms. As proof-of-principle, we further dissect the interplay between one functional hit, GSPT1, with EBOV polymerase. Our results suggest that at early times during EBOV infection GSPT1 exhibited antiviral activity, but at later times on infection EBOV hijacked GSPT1 to facilitate the termination step of viral transcription. Likewise, our data suggest that EBOV can counteract UPF1-restriction documented for several viruses, and EBOV employs UPF1 to positively influence viral replication. Previously documented studies aimed at elucidating EBOV protein-host interactomes typically used cells overexpressing a single viral bait protein. Such bait proteins and protein-protein interactions analyzed using traditional affinity-purification mass spectrometry (AP-MS) experiments must be stable to facilitate affinity-purification and analysis, which could miss. However, the EBOV polymerase can be unstable and its interactions with other cellular or viral factors can be dynamic. To capture both stable and dynamic protein interactions, we used a proximity-labeling approach via a split-TurboID tagged-EBOV polymerase and cells expressing a functional EBOV MG that recapitulates EBOV RNA synthesis events. This system also allowed us to compare the cellular interactome of EBOV polymerase with and without VP30, an essential viral factor to regulate EBOV transcription but not replication.

In the absence of VP30, we identified eight functional hits that interacted with EBOV polymerase. Of these, ZC3H7A, HNRNPA1, and MRE11 which have been implicated in SARS-CoV2 (Hoffmann et al., 2021), Hepatitis C virus (Kim et al., 2007; Rios-Marco et al., 2016), and Herpes simplex virus infection (Kondo et al., 2013), respectively. These host factors also suppressed EBOV infection, at least at early times of infection, consistent with the existence of conserved host restriction factors that may respond to infection by different viruses. We also identified the dsRNA sensor EIF2AK2/PKR, known to be antagonized by EBOV VP35 (Feng et al., 2007; Schumann et al., 2009).

In the presence of VP30, we found 19 functional hits within the EBOV-polymerase interactome. These hits are functionally enriched in four biological processes: cytoskeleton organization (DOCK7, CTTN, CORO7, CNN3, AMOT, FLNA), chaperone-mediated protein folding (UNC45A and CCT8), negative regulation of translation (EIF4E2 and GIGYF2) and RNA catabolic processes (UPF1 and GSPT1). These biological processes may contribute to EBOV RNA synthesis as the formation of cytoplasmic viral inclusion bodies may involve rearrangement of the cytoskeleton; the EBOV polymerase folding may need host chaperones; the production of EBOV proteins requires host translational machinery; and newly synthesized viral RNAs may be subjected to cellular mRNA decay pathway.

Two EBOV polymerase interactors, UPF1 and GSPT1, are critical players in the cellular nonsense-mediated decay (NMD) pathway. To trigger the NMD pathway, UPF1 recognizes mRNA targets marked with aberrant features, such as an unusually long 3’UTR or an upstream open-reading frame (uORF) linked to the presence of a premature-stop codon. This process is typically coupled with translation termination and promoted by the interaction between UPF1 and the termination complex (GSPT1/eRF3-eRF1) (Kurosaki et al., 2019). The NMD pathway has been shown to restrict multiple positive- and double-strand RNA viruses, but not retroviruses or influenza virus (Ajamian et al., 2008b; Balistreri et al., 2017; Declercq et al., 2020; Popp et al., 2020; Tran et al., 2021). How NMD affects negative-strand RNA viruses like EBOV that replicate in the cytoplasm remains to be determined. A uORF is present in 4/7 EBOV mRNAs (Shabman et al., 2013), presumably qualifying them as NMD substrates. However, our results revealed that UPF1 and GSPT1 exerted only a brief restriction of EBOV infection at the onset of viral infection, similar to that seen for influenza virus infection (Tran et al., 2021). At later times of infection, neither UPF1 nor GSPT1-depletion enhanced EBOV multiplication in human hepatocytes.

EBOV instead hijacked NMD factors to its own benefit. For instance, our data indicated that GSPT1 interacts with EBOV polymerase and promotes viral transcription. As with other mononegaviruses including vesicular stomatitis virus (VSV) (Barr et al., 2002) and respiratory syncytial virus (RSV) (Fearns and Collins, 1999), EBOV transcription is highly regulated by *cis*-acting elements, which comprise nearly 30% of the viral genome (Biedenkopf et al., 2016; Brauburger et al., 2014; Muhlberger et al., 1999; Weik et al., 2002). Studies examining EBOV transcription regulation have largely focused on the virus-encoded, *trans*-acting factor VP30 and its role in activating EBOV transcription initiation. Our results revealed an additional mechanism of transcriptional regulation where the host factor GSPT1 modulates EBOV transcription termination. Given the decreased level of viral genomic RNA observed in GSPT1-KD cells, we cannot rule out a possible separate role for GSPT1 in facilitating EBOV genome replication.

Pharmacological targeting of GSPT1 protein via CC-90009 correlated with strong reduction in viral RNA synthesis and production of infectious progeny. Downstream physiological effects of CC-90009 treatment (Surka et al., 2021) may synergistically contribute to EBOV inhibition. Future studies should dissect the exact mechanisms of CC-90009-mediated anti-EBOV activity and determine whether CC-90009 can similarly inhibit other filoviruses. CC-90009 is a drug candidate currently in Phase 1b clinical trial to treat patients with acute myeloid leukemia. Hence, our results illuminate a possible path to repurpose CC-90009 as an anti-EBOV agent.

Our results also imply that UPF1 plays a supporting role in EBOV RNA synthesis, a finding similar to that reported for HIV (Ajamian et al., 2008a; Serquina et al., 2013). Future studies should verify the interaction between UPF1 and EBOV polymerase and explore the potential interaction between UPF1 and EBOV RNAs. Staufen-mediated mRNA decay (SMD), a quality-control mechanism related to NMD that acts in parallel (Park and Maquat, 2013) does not restrict EBOV infection either. EBOV rather recruits the Staufen1 protein to support viral RNA synthesis (Fang et al., 2018). Our study highlights the ability of EBOV to not only counteract cellular mRNA decay mechanisms, but to also capitalize cellular mRNA decay factors to enhance or to help regulating viral replication and gene expression.

## LIMITATIONS OF THE STUDY

The EBOV MG system we used in our proximity proteomic experiment is active in the absence of EBOV VP30. This feature allowed us to probe the differential EBOV polymerase interactome in the presence or absence of VP30. VP30 is indispensable during authentic EBOV infection. Thus, we only examined high-confidence hits revealed in our interactomes that were confirmed in our siRNA-based functional screen with authentic EBOV infection. We also validated the protein-protein interaction between EBOV polymerase and GSPT1 using complementary biochemical and imaging approaches.

It should be noted that our siRNA-based functional screen only captured a snapshot of EBOV infection at 48 hours post-infection at a chosen MOI. Thus, the assigned pro- or antiviral role of a given hit is specific to the infection parameters and may not apply across the entire time course of EBOV infection. As seen for GSPT1 and UPF1, the same host factor can be antiviral very early in infection but can be proviral later. Our results have provided the basis for comprehensive follow-up studies to determine the roles of each functional EBOV polymerase interactor in the context of the complete EBOV life cycle in relevant cell types and appropriate animal models.

The EBOV MG system being used models the activity of a monocistronic EBOV genome and precludes transcription readthrough events. Future experiments using a tetracistronic EBOV MG system (Schmidt et al., 2018) should verify the supporting role for GSPT1 in EBOV transcription termination described in the current study. Lastly, alternative splicing results in multiple isoforms of GSPT1 protein that share a common C terminus but have different N termini. The level of GSPT1 proteins and GSPT1 transcripts analyzed here was taken as an overall representation of all isoforms assuming that they are interchangeable. Whether a specific isoform of GSPT1 is responsible for the phenotype we observed in EBOV infection should be determined, as well as any functional consequence of the GSPT1 fragment manifested in EBOV infected cells.

## ACKNOWLEDGMENTS

We thank Jonathan Towner (CDC) and Stuart Nichol (CDC) for providing the EBOV full-length clone, Yoshihiro Kawaoka (University of Wisconsin) for providing the pCEZ-NP, VP35, L, VP30, and the pHH21-3E5E-Fluc plasmids, as well as sharing the anti-VP30 antibody. We thank Beatrice Cubitt from the de la Torre Lab (Scripps Research, CA) for helping with cloning of pCI-FLAG-GSPT1 plasmid. We thank Diptiben Parekh (LJI) for plasmid preparations, Sharon Schendel (LJI) for manuscript editing, Zbigniew Mikulski of the Microscopy Core Facility (LJI) for microscopy training, and NIH S10OD021831 for sponsoring the Zeiss LSM 880 microscope. This research was supported by institutional funds of La Jolla Institute for Immunology. J.F. was supported by the Donald E. and Delia B. Baxter Foundation Fellowship.

## AUTHOR CONTRIBUTIONS

J.F., J.C.T., and E.O.S. designed research and wrote the paper; J.F. performed all non-BSL4 experiments and analyzed data; C.P. performed all BSL-4 experiments; A.B. provided supervision for BSL-4 research; G.T. and G.C performed proteomic analysis; K.F.C. and A.Y.T provided a critical resource; all authors edited and approved the paper.

## DECLARATION OF INTERESTS

The authors declare no competing interest.

## STAR METHODs

### KEY RESOURCES TABLE

| REAGENT or RESOURCE | SOURCE | IDENTIFIER |
| --- | --- | --- |
| **Antibodies** |  |  |
| Streptavidin-HRP | Thermofisher, Invitrogen | Cat#SA10001 |
| Mouse, anti-HA, 16B12 (monoclonal) | Biolegend<br>Cat#901513 | Cat#901513 |
| Human, anti-EBOV NP, KZ51 (monoclonal) | Absolute antibody | Cat#Ab00692-10.0 |
| Rabbit, anti-VP30 (#1-1) | A kind gift from Peter Halfmann and Yoshihiro Kawaoka | N/A |
| Mouse, anti-EBOV VP24_21.5.2.5 (monoclonal) | A kind gift from Peter Halfmann and Yoshihiro Kawaoka | N/A |
| Mouse, anti-EBOV VP40 | Customized antibody by Zalgen labs/<br>Autoimmune technology | N/A |
| Human, BDBV223 (Cross-reactive) Anti-EBOV GP2 | A kind gift from James E. Crowe (Flyak et al., 2018) | N/A |
| Rabbit, polyclonal anti-EBOV L | IBT Bioservices | Cat#0301-045 |
| Rabbit, polyclonal anti-GSPT1/eRF3 | Abcam | Cat#ab49878 |
| Rabbit, anti-UPF1 (D15G6) (monoclonal) | Cell signaling technology | Cat#12040 |
| Streptavidin-AlexaFluor488 | Thermofisher/Invitrogen | Cat#s11223 |
| Goat, anti-mouse-HRP | Thermofisher/Invitrogen | Cat#31437 |
| Goat, anti-rabbit-HRP | Southern Biotech | Cat#4050-05 |
| Goat, anti-human-HRP | Thermofisher/Invitrogen | Cat#A18811 |
| Goat, anti-mouse-AlexaFluor568 | Thermofisher/Invitrogen | Cat#A11004 |
| Goat, anti-mouse-AlexaFluor647 | Thermofisher/Invitrogen | Cat#A21236 |
| Goat, anti-rabbit-AlexaFluor488 | Thermofisher/Invitrogen | Cat#A11008 |
| Goat, anti-rabbit-AlexaFluor647 | Thermofisher/Invitrogen | Cat#A27040 |
| Streptavidin-magnetic beads | ThermoFisher | Cat#88817 |
| Anti-FLAG M2 affinity Gel | Sigma-Aldrich | Cat#A2220-5ML |
| **Bacterial and virus strains** |  |  |
| EBOV-eGFP (strain Mayinga) | Bukreyev Lab | Initially provided by Jonathan Towner (CDC) and Stuart Nichol (CDC) (Towner et al., 2005) |
| **Chemicals, peptides, and recombinant proteins** |  |  |
| Biotin | Sigma-Aldrich | Cat#B4501 |
| CC-90009 (>98%) | MedKoo Biosciences | Cat# 207005, Lot# S20S01C02 |
| Dimethyl Sulfoxide (DMSO) | Fisher BioReagents | Cat# BP231-100 |
| Lipofectamine™ RNAiMAX Transfection Reagent | ThermoFisher/Invitrogen | Cat#13778150 |
| TransIT®-LT1 Transfection Reagent | Mirus | Cat#MIR2306 |
| Sequencing Grade Modified Trypsin | Promega | Cat#V5111 |
| NEBBuilder HiFi DNA Assembly Master Mix | NEB | Cat# E2621 |
| In-Fusion® HD Cloning Plus | Takara | Cat# 638920 |
| 16% Paraformaldehyde Aqueous Solution, EM Grade, Ampoule 10ML | Electron Microscopy Sciences | Cat#15710 |
| ProLong™ Gold Antifade Mountant | ThermoFisher/Invitrogen | Cat#P36934 |
| Normal Goat Serum | ThermoFisher/Invitrogen | Cat#PCN5000 |
| Hoechst 33342, Trihydrochloride, Trihydrate, 100 mg | ThermoFisher/Invitrogen | Cat#H1399 |
| RIPA Buffer (10X) | Cell Signaling Technology | Cat#9806S |
| cOmplete™, EDTA-free Protease Inhibitor Cocktail | Roche | SKU# 11873580001 |
| Benzonase® Nuclease, Purity > 90% | Millipore/Sigma | Cat#70746 |
| NuPage Sample reducing agent (10X) | ThermoFisher | Cat#NP0009 |
| NuPage LDS sample buffer (4X) (Laemmli) | ThermoFisher | Cat#NP0008 |
| TRIzol™ Reagent | ThermoFisher/Invitrogen | Cat# 15596018 |
| DEPC-Treated Water | ThermoFisher/Invitrogen | Cat#AM9920 |
| Iodoacetamide (IAA) | Sigma-Aldrich | SKU#I1149-5G |
| **Critical commercial assays** |  |  |
| Pierce™ BCA Protein Assay Kit | Thermo Scientific | Cat#23225 |
| Pierce™ Silver Stain for Mass Spectrometry | Thermo Scientific | Cat#24600 |
| SuperSignal™ West Pico PLUS Chemiluminescent Substrate | Thermo Scientific | Cat#34578 |
| Luciferase Assay System | Promega | Cat#E4550 |
| CellTiter-Glo® 2.0 Cell Viability Assay | Promega | Cat#G9241 |
| High-Capacity cDNA Reverse Transcription Kit with RNase Inhibitor | ThermoFisher scientific (Applied Biosystems) | Cat#4374966 |
| SYBR™ Select Master Mix | ThermoFisher scientific (Applied Biosystems) | Cat#4472908 |
| Direct-zol RNA Miniprep Plus | Zymo research | Cat#R2071 |
| ZipTip with 0.2 µL C18 resin | Millipore | Cat# ZTC18M096 |
| TMT10plex™ Isobaric Label Reagent Set, 1 x 0.8 mg | Thermo Scientific | Cat# 90110 |
| **Deposited data** |  |  |
| Raw proximity proteomic data of EBOV polymerase interactome | This study | Table S1 |
| **Experimental models: Cell lines** |  |  |
| HEK 293T | ATCC | Cat#CRL-3216 |
| VERO C1008 [Vero 76, clone E6, Vero E6] | ATCC | Cat# CRL-1586 |
| Huh7 used in BSL4 | A kind gift from Mariano Garcia-Blanco |  |
| Huh7 used in BSL2 | A kind gift from Luke Wiseman |  |
| **Oligonucleotides** |  |  |
| Primers for RT-qPCRs, see Table S3 | This paper | N/A |
| siRNAs for knockdowns, see Table S2 | This paper | N/A |
| AllStars Negative Control siRNA | Qiagen | Cat#1027281 |
| Oligo dT(20) Primer | Thermofisher/invitrogen | Cat#18418020 |
| **Recombinant DNA** |  |  |
| Plasmid: pCEZ-NP | (Jasenosky et al., 2010b) | N/A |
| Plasmid: pCEZ-VP35 | (Jasenosky et al., 2010b) | N/A |
| Plasmid: pCDNA5-VP35-HA | (Fang et al., 2018) | N/A |
| Plasmid: pCAGGS-VP35-HA | This study | N/A |
| Plasmid: pCAGGS-L | A kind gift from Thomas Hoenen | N/A |
| Plasmid: pCAGGS-VP35-sNTurboID | This study | N/A |
| Plasmid: pCEZ-VP30 | (Jasenosky et al., 2010b) | N/A |
| Plasmid: pCEZ-L | (Jasenosky et al., 2010b) | N/A |
| Plasmid: pCEZ-L-sCTurboID | This study | N/A |
| Plasmid: pHH21-3E5E-Fluc | (Jasenosky et al., 2010b) | N/A |
| Plasmid: pCI-empty | This study | N/A |
| Plasmid: pCI-MS2V5-eRF3a F76a | (Fatscher et al., 2014) | Plasmid #65811 |
| Plasmid: pCI-FLAG-GSPT1 | This study | N/A |
| Plasmid: pCAGGS-eGFP | This study | N/A |
| Plasmid: pCAGGS-empty | This study | N/A |
| Plasmid: pLX304 CMV HA-HaloTag-FRB-sTurboID (C) | (Cho et al., 2020) | Plasmid: #153003 |
| Plasmid: pLX304 CMV FKBP-V5-sTurboID (N) | (Cho et al., 2020) | Plasmid: #153002 |
| Plasmid: pFastBac™ Dual Expression Vector | ThermoFisher/Gibco | Cat#10712024 |
| **Software and algorithms** |  |  |
| GraphPad Prism 9 | GraphPad | N/A |
| Cytoscape v3.8.2 | Cytoscape | https://cytoscape.org/ |
| Image Lab Software | Bio-Rad | https://www.bio-rad.com/en-us/product/image-lab-software |
| SparkControl™ | Tecan |  |
| BioRender | BioRender | https://biorender.com/ |
| Thermo Scientific™ HCS Studio Cell Analysis Software | Thermo Scientific | CX51110 |
| ZEISS ZEN 3.4 Blue edition | ZEISS | https://www.zeiss.com/ |
| FIJI | (Schindelin et al., 2012) | https://imagej.net/software/fiji/ |
| ImageJ | (Schneider et al., 2012) | https://imagej.nih.gov/ij/ |
| SnapGene Viewer | SnapGene | https://www.snapgene.com/snapgene-viewer/ |
| STRING | (Szklarczyk et al., 2019) | https://string-db.org/ |
| Proteome Discover v 2.3 | Thermo Scientific |  |
| SEQUEST | Yates Laboratory | http://proteomicswiki.com/wiki/index.php/SEQUEST |
| Scaffold 4 | Proteome Software |  |

## RESOURCE AVAILABILITY

### Lead contact

Further information and requests for resources and reagents should be directed to and will be fulfilled by the Lead Contact, Erica Ollmann Saphire.

### Materials availability

All requests for unique reagents generated in this study should be directed to and will be fulfilled by the Lead Contact author. Materials will be made available through the authors upon execution of a Material Transfer Agreement.

### Data and code availability

Raw proteomic data reported in this study are reported in **Table S1**.

## EXPERIMENTAL MODEL AND SUBJECT DETAILS

### Cell cultures

Human hepatocytes Huh7 and human embryonic kidney cells HEK 293T were maintained in Dulbecco’s modified Eagle medium (DMEM-GlutaMAX) supplemented with 4.5 g/L D-Glucose, 10% fetal bovine serum (FBS), penicillin (100 U/ml), streptomycin (100 μg/ml). For cells used in viral infection experiment, Huh7 were cultured in Dulbecco’s modified Eagle medium (DMEM) supplemented with 10% FBS and 50 μg/ml Gentamicin Sulfate; Vero-E6 cells were cultured in Minimum Essential Medium (MEM) supplemented with 10% FBS, 1% MEM Non-essential Amino Acid Solution, 1% Sodium pyruvate solution, and 50 μg/ml Gentamicin Sulfate. All cells were grown at 37°C and 5% CO2.

### Virus stock

EBOV strain Mayinga expressing Emerald green fluorescent protein (eGFP) was propagated in Vero-E6 cells with 500,000 plaque-forming unit (PFU) per T225 flask (at 90% confluency). EBOV-eGFP was harvested 5 days post-infection. All experiments using infectious EBOV were performed under biosafety level 4 (BSL-4) conditions at the Galveston National Laboratory. Virus inactivation was performed according to standard operating procedures.

## METHOD DETAILS

### Plasmids and cloning

To generate pCEZ-L-sCTurboID, a fragment of L coding sequence was first subcloned into a pFastbacDual intermediate plasmid using the natural restriction sites Pac1 and Hpa1. From there, a fragment of sequence encoding the sC-TurboID and a FLAG-tag sequence, flanked by two flexible linkers was internally inserted at the position 1705/1706 (TTIP/Q). This insertion was PCR amplified from pLX304 CMV HA-HaloTag-FRB-sTurboID (C), and inserted to the intermediate plasmid using NEBuilder HiFi DNA Assembly Master Mix (NEB). The L-sNTurboID fragment was ligated back to the pCEZ-L backbone using the same restriction sites Pac1 and Hpa1 by NEB T4 ligase.

To generate pCAGGS-VP35-HA, the coding sequence of VP35-HA was subcloned to the plasmid pCAGGS-empty using the restriction sites Kpn1 and Xho1 by standard ligation method. To generate pCAGGS-VP35-sNTurboID, pcDNA5-VP35-HA (gift from Mariano Garcia-Blanco) was linearized. The fragment of sNTurboID was PCR amplified from pLX304 CMV FKBP-V5-sTurboID (N) and cloned into the pcDNA5-VP35-HA using NEBuilder HiFi DNA Assembly Master Mix (NEB). The fragment of VP35-sNTurboID, including the C-terminal HA tag, was subcloned to pCAGGS plasmid using restriction sites Kpn1 and Nhe1 and standard ligation method.

To generate pCI-empty, the coding sequence of MS2V5-eRF3a-F76a was removed from pCI-MS2V5-eRF3a F76a plasmid (a gift from Niels Gebring) by PCR and re-circularized using NEBuilder HiFi DNA Assembly Master Mix (NEB). To generate pCI-FLAG-GSPT1, the coding sequence of MS2V5 was removed from pCI-MS2V5-eRF3a F76a plasmid and an N-terminal FLAG tag was inserted to the same plasmid by PCR and re-circularized using NEBuilder HiFi DNA Assembly Master Mix (NEB). The F76a mutation was mutated back to wildtype using In-fusion cloning (Clontech).

The sequence of all engineered plasmid DNA was confirmed by Sanger sequencing.

### EBOV minigenome assay

HEK 293T cells (2 × 10^5^ per well) were seeded in 24-well plates one day prior to transfection with the following plasmids: 100 ng pCEZ-NP, 100 ng pCEZ-VP35, 75 ng pCEZ-VP30, 750 ng pCEZ-L, 62.5 ng EBOV minigenome encoding a firefly luciferase reporter (pEE21). In some cases: pCEZ-VP30 was replaced with equal mass of control pCAGGS-empty plasmid; pCEZ-L and/or pCEZ-VP35 were replaced with a tagged variant. A negative control was included in every assay in which the pCEZ-L plasmid was replaced with equal mass of control pCAGGS-empty plasmid to verify minigenome activity mediated by EBOV polymerase. Plasmids were transfected with TransIT-LT1 transfection reagent (3 µl/µg plasmid). At 48 hours post-transfection, cells were lysed and subjected to the luciferase reporter assay (Promega).

To measure the effect of GSPT1 knockdown on EBOV minigenome activity, we modified the assay as below. HEK 293T cells (1 × 10^5^ per well) were seeded in 24-well plates one day prior to siRNA or plasmid. To knockdown GSPT1, cells were transfected with 10 nM of the siRNA, GSPT1si8, using Lipofectamine RNAiMAX. In control wells, cells were transfected with 10 nM of the Allstar negative control siRNA (NSC) using Lipofectamine RNAiMAX. To overexpress GSPT1, cells were transfected with 100/200/400 ng of pCI-FLAG-GSPT1 (per well) using TransIT-LT1. In control wells, cells were transfected with 200 ng of pCI-empty control vector using TransIT-LT1. At 24 hours post-transfection, cells were transfected with the EBOV MG system described earlier. In the control plate, cells were transfected with 100 ng of pCAGGS-eGFP (per well) using TransIT-LT1 to evaluate the effect of GSPT1 modulation on a control reporter expression. Two days after the second transfection, cell lysate was harvested in 1X passive lysis buffer for luciferase reporter assay or in GFP lysis buffer (50mM Tris pH7.4, 1mM EDTA, 0.5% NP-40, 150mM NaCl) to quantify the fluorescence intensity. A fraction of each lysate was used to quantify the total protein by BCA assay. Raw luciferase intensity or GFP intensity of each sample was first normalized to the BCA quantified protein amount. The resulting relative luciferase reporter activity or GFP reporter activity was further normalized to that of the control: NSC for the GSPT1 knockdown experiment, pCI-empty for the GSPT1 overexpression experiment.

### Biotinylation with polymerase-TurboID fusion protein

HEK 293T cells seeded in 24-well plates were transfected with EBOV minigenome system including the wild-type polymerase or the polymerase-TurboID fusion. Biotin (500 μM) was added to cell culture media 2-3 days post-transfection. Multiple labeling time points were sampled for optimization. The reaction was stopped by placing cells on ice, the excess biotin was removed by DPBS washes, and soluble whole cell lysates were collected for western blot analysis using streptavidin-HRP.

### Immunofluorescent analysis (IFA) using confocal microscopy

Acetone cleaned, glass coverslips (1.5 mm thickness) placed inside 24-wells were treated with human fibronectin (50 mg/ml) for 20 min at 37 ºC incubator, prior to HEK 293T cells seeding (4×10^4^/well). Twenty-four hours later, the monolayer was transfected with the EBOV minigenome system using Trans-IT LT1 as described earlier. For in-cell biotinylation experiment: at the same day of transfection, biotin (500 μM) was added to the culture media allowing for 18 hours in-cell biotinylation in EBOV minigenome transfected cells. Labeling was stopped by washing cells with cold DPBS for three times. For all IFA specimens, cells were fixed in 4% paraformaldehyde for 15 mins, quenched with 20mM Glycine-DPBS for 5 mins, permeabilized with 0.1% Triton-X100-DPBS for 10 mins, and blocked in 1% normal goat serum for at least one hour, all at room temperature. Primary antibody incubations were performed at room temperature for two hours or at 4 ºC overnight. Specimens were washed three times with PBS-0.1% Tween-20 and incubated with secondary antibodies for one hour at room temperature. Nuclei were counterstained by Hoechst. Confocal images were acquired on Zeiss LSM880-airyscan system under super-resolution mode, using a 63x/NA1.4 oil objective.

For samples shown in the Figure 1D, mouse anti-HA (1:500) and rabbit anti-VP30 (1:500) antibody were used as the primary antibodies, goat anti-mouse AlexaFluor647 (1:500) and goat anti-rabbit AlexaFluor 568 (1:500) were used as the secondary antibodies. Together these combinations were used to probe HA-tagged EBOV VP35 (VP35-HA and VP35-sNTurboID) and the wildtype VP30. Streptavidin-AlexaFluor488 (1:500) was added along with secondary antibodies to probe biotinylated proteins.

For samples shown in Figure 5B, mouse anti-HA (1:500) and rabbit anti-GSPT1 (1:500) antibody were used as the primary antibodies, goat anti-mouse AlexaFluor568 (1:500) and goat anti-rabbit AlexaFluor 488 (1:500) were used as the secondary antibodies. Together these combinations were used to probe EBOV VP35-HA and the endogenous GSPT1 protein.

### Sample preparation for in-cell biotinylation and streptavidin enrichment

Protocol adapted from a previous study (Branon et al., 2018). HEK 293T cells were seeded in 6-wells plate (1×10^6^ cells per well) and transiently transfected using Trans-IT LT1 a day after, with EBOV MG plasmids including plasmid expressing wild-type or the TurboID-fused viral polymerase. To enable in-cell biotinylation, at two days post-transfection, monolayers expressing EBOV MG are incubated with 500 μM biotin containing complete media for indicated times. Biotinlyation was stopped by washing monolayers with cold DPBS twice before collecting the cell pellets by centrifugation at 1500 rpm for 3 mins at 4 ºC. Cell pellets (1V) were lysed in 1V of 1X RIPA buffer containing cOmplete™, EDTA-free Protease Inhibitor Cocktail (Roche) and Benzonase® Nuclease (Millipore) followed by incubation on ice for 15-20 mins. Soluble lysates were clarified by centrifugation at 13000 rpm for 5 mins at 4 ºC.

For streptavidin enrichment, streptavidin-magnetic beads (80 μl slurry/mg of total protein) were washed with 1X RIPA buffer twice and incubated with 2-3 mg of soluble cell lysate for one hour at room temperature. The beads were subsequently washed twice with 1 ml of 1X RIPA buffer, once with 1 ml of 1M KCL, once with 1 ml of 0.1M Na_2_CO_3_, once with 1 ml of 2M Urea in 10 mM Tris-HCl (pH 8.0), and with twice with 1 ml 1X RIPA buffer. Washed beads were then either eluted in 50 uls of NuPage LDS sample buffer (4X) supplemented with 20 mM DTT and 2 mM biotin for quality check or processed to on beads digestion. Quality checks for successful streptavidin enrichment include silver staining and western blotting of SDS-PAGE-separated input and streptavidin-enriched samples side-by-side.

For on-bead digestion, proteins bound to beads were further washed twice with 50 mM Tris HCL (pH 7.5) and two washes of 2 M urea/50 mM Tris buffer (pH 7.5). A final volume of 80 μl of 2 M urea/50 mM Tris containing 1 mM DTT and 0.4 μg trypsin was added to washed beads for overnight digestion at 37 ºC shaker. Digested supernatants containing biotinylated peptides were transferred to a new tube. The streptavidin beads were washed twice with 60 μl of 2 M urea/50 mM Tris buffer (pH 7.5) and the rests were combined with the on-bead digest supernatant. The eluate was reduced with 4 mM DTT for 30 mins at room temperature with shaking, alkylated with 10 mM IAA (iodoacetamide) for 45 mins in the dark at 25 ºC with shaking. An additional 0.5 μg of trypsin was added to the sample and the digestion was completed overnight at 37 ºC shaker. After final digestion, samples were acidified by adding formic acid to a final concentration of 1 % and stored in - 80 ºC freezer before sending to Scripps Research (Florida) Proteomic core for further processing, TMT labeling, and mass-spectrometry analysis.

### Proximity proteomics (with tandem-mass-tag/TMT labeling)

Following on-bead trypsin digestion of biotinylated proteins, LASSA (WT and TurboID) and EBOLA (WT, TurboID MINUS and TurboID PLUS) samples were acidified by the addition of 1% formic acid and desalted with 2 μg-capacity C18 ZipTips, respectively, and dried using vacuum. Peptides were resuspended in 100mM TEAB and labelled with TMT labels (10-plex, with one TMT label not used) according to the manufacturer’s instructions and pooled.

The labelling scheme for three biological replicates of each condition was as follows:

|  |  |  |  |
| --- | --- | --- | --- |
| EBOLA sample | WT_pol+VP30_01 | WT_pol+VP30_02 | WT_pol+VP30_03 |
| TMT<br>10-plex label | 126 | 127C | 128C |
| EBOLA sample | TurboID_pol-VP30_01 | TurboID_pol-VP30_02 | TurboID_pol-VP30_03 |
| TMT<br>10-plex label | 127N | 128N | 131 |
| EBOLA sample | TurboID_pol+VP30_01 | TurboID_pol+VP30_02 | TurboID_pol+VP30_03 |
| TMT<br>10-plex label | 129N | 130N | 129N |

The pooled and multiplexed samples were dried under vacuum, re-solubilized in 1% TFA and then desalted using 2 μg-capacity C18 ZipTips (Millipore, Billerica, MA), and then dried once again using vacuum. For mass spectrometry, dried TMT-labelled peptides were reconstituted in 5 μL of 0.1% TFA, vortexed briefly, and then sonicated for 15 mins. The peptides were subsequently on-line eluted into a Fusion Tribrid mass spectrometer (ThermoFisher Scientific, San Jose, CA) from an Acclaim PepMapTM RSLC C18 nano Viper analytical column (2 mm, 100 Å, 75-μm ID × 50 cm, Thermo Scientific, San Jose, CA) using a gradient of 5-25% solvent B (80/20 acetonitrile/water, 0.1% formic acid) in 180 mins, followed by 25-44% solvent B in 60 mins, 44-80% solvent B in 0.1 min, a 5-mins hold of 80% solvent B, a return to 5% solvent B in 0.1 min, and finally a 20-mins hold of solvent B. All flow rates were 300 nl/min delivered using a nEasy-LC1000 nano liquid chromatography system (Thermo Fisher Scientific, San Jose, CA). Solvent A consisted of water and 0.1% formic acid. Ions were created at 2.1-2.5 kV using the EASY-SprayTM ion source held at 50 ºC (Thermo Fisher Scientific, San Jose, CA). A synchronous precursor selection (SPS)-MS3 mass spectrometry method was used based on published protocol (Ting et al., 2011), scanning between 380-2000 m/z at a resolution of 120,000 for MS1 in the Orbitrap mass analyzer, and performing CID at top speed in the linear ion trap of peptide monoisotopic ions with charge 2-8, using a quadrupole isolation of 0.7 m/z and a CID energy of 35%. The top 10 MS2 ions in the ion trap between 400-1200 m/z were then chosen for HCD at 65% energy and detection in the Orbitrap at a resolution of 60,000 and an AGC target of 1E5 and an injection time of 120 msec (MS3). Three technical replicates were performed for each sample.

### Bioinformatic analysis of mass spectrometry data

The raw data technical replicates were added as “fractions” into Proteome Discover v 2.3 (Thermo Fisher Scientific, San Jose, CA). As it is the default option for this software, peptide spectrum match raw values of both technical replicates from the same biological sample were summed for the same peptide group discovery. Quantitative analysis of the TMT experiments was performed simultaneously to protein identification. The precursor and fragment ion mass tolerances were set to 10 ppm, 0.6 Da, respectively), enzyme was Trypsin with a maximum of 2 missed cleavages. Steptavidin FASTA and Uniprot Human proteome FASTA file with added sequences specific for EBOV proteins and TurboID fusion proteins, were used in SEQUEST searches. The impurity correction factors obtained from Thermo Fisher Scientific for each kit was included in the search and quantification. The following settings were used to search the streptavidin enriched data; dynamic modifications; Oxidation / +15.995Da (M), Deamidated / +0.984 Da (N, Q), TMT6plex / +229.163 Da (K), Biotin / +226.078 Da (K) and static modifications of TMT6plex / +229.163 Da (N-Terminus), Carbamidomethyl +57.021 (C). Only unique+ Razor peptides were considered for quantification purposes. Target Decoy feature of Proteome Discoverer 2.3 was used to set a false discovery rate (FDR) of 0.01. Total peptides quantified were normalized to *Streptavidin* to adjust for loading bias and Protein Abundance Based method was used to calculate the protein level ratios. Low Abundance Resampling method was used to impute missing data and co-isolation threshold and SPS Mass Matches threshold were set to 50 and 65, respectively. ANOVA (Individual Proteins) was performed using Proteome Discoverer 2.3 workflow and FDR of 0.05 was chosen as the cut off to identify the top tier proteins that are differentially expressed across WT and TurboID samples.

Sequences added to the Uniprot Human proteome database for this experiment:

- Ebola VP30
- Ebola NP
- Ebola VP35_wildtype
- Ebola L_wildtype
- Ebola VP35_sCTurboID fusion protein
- Ebola L_sNTurboID fusion protein

### Thresholding and network analysis of viral polymerase-proximity interactome

To curate a list of high confident hits for both viral polymerase-proximity interactome, we applied a threshold of > 1 for log_2_ transformed relative (rel.) abundance ratio (TurboID_pol/WT_pol) and a threshold of < 0.05 for the adjusted P value for each abundance ratio. To analyze the differential interactome of EBOV polymerase with and without VP30, since the experimental set up only included the wild-type EBOV polymerase with VP30 as the control, abundance ratios of proteins identified in the TurboID polymerase interactome with or without VP30 were first normalized to the WT pol +VP30 control to get the log_2_(fold change) value. Subsequently, the difference of the two log_2_(fold change) values, Δlog_2_(fold change), was determined as a measure of the differential interactome of EBOV polymerase with and without VP30. To curate a list of high confidence hits, we retained the same threshold of “< 0.05” for the adjusted P value for corresponding abundance ratio of (TurboID_pol -VP30/WT_pol) and further applied a threshold of Δlog_2_(fold change) >1. Above mentioned lists including critical parameters used in thresholding were presented in **Table S1**.

High-confidence hits in both EBOV polymerase interactomes in the presence and absence of VP30 were displayed as protein-protein networks using Cytoscape. The web-based bioinformatic analyzer STRING was used to perform functional enrichment analysis to cluster nodes involved in the same biological process (FDR< 1%).

### siRNA screen with authentic EBOV infected cells

Huh7 cells (5 × 10^3^ per well) were seeded in clear-bottom, black 96-wells plate, leaving edge wells filled with DPBS. Twenty-four hours, Huh7 monolayers were transfected with 10 nM of individual siRNAs using Lipofectamine RNAiMAX (0.1 ul per well). Knock down (KD) of each target was performed using four individual siRNA transfections, each one performed in triplicate. Each plate includes two sets of triplicated internal control wells transfected with 10 nM AllStar negative control (NSC) to account for plate-to-plate variations. Forty-eight hours later, plates were transferred to BSL4 for infection with EBOV-eGFP at indicated MOIs. Infected monolayers were fixed and inactivated by 10% formalin at two days post-infection and removed from BSL4 facility. Fixed monolayers were stained with Hoechst (1 µg/ml in DPBS) for 15 mins at room temperature prior to imaging. Percentage of EBOV-eGFP infected Huh7 cells (% infection) and cell count with different siRNA treatment were quantified by CellInsight™ CX5 High Content Screening (HCS) Platform (ThermoFisher) using a 10x objective.

Relative % virus infection and relative cell count were calculated by normalizing raw percent viral infection rate and cell count of individual siRNA treatment to corresponding control wells in each plate. Heat maps showing relative %virus infection and relative cell count were generated by GraphPad Prism 9 using the normalized value of each individual siRNA treatment. The mean of numerical values from each triplicated well are used for plotting. To quantitatively analyze the impact of siRNA treatment on viral infection and cell count, multiple t-tests were performed, comparing normalized % infection of each siRNA treatment to corresponding NSC control. Bubble plots were generated by plotting numerical values of normalized cell count of each siRNA treatment against their value of normalized % infection if their P value (represented by the size of each data point) was < 0.05. Outliers with cell count significantly different from the average by one standard deviation were ruler out because the effect of siRNA treatment on %infection in these cases can be confounded by a marked alteration in the total number of cells. siRNAs used for the screen were from the Genome-wide ON TARGET-Plus (OTP) Human siRNA library from Dharmacon/Thermo Scientific. Sequence information of each siRNA been used is curated in **Table S2**.

### Co-Immunoprecipitations (co-IP)

Co-IP reactions were performed as previously described (Fang et al., 2018). HEK 293T cells (1×10^6^) were seeded in 6-wells plate. Twenty-four hours later, cells in each well were transfected with 500 ng of pCI-empty or pCI-FLAG-GSPT1 plasmid combined with 3 μg of pCAGGS-L and 400 ng of pCAGGS-VP35, plus and minus 300 ng of pCEZ-VP30, using TransIT LT1 transfection reagent. For experiments probing interactions between GSPT1 and EBOV VP35 or VP30, 1 μg of pCI-empty or pCI-FLAG-GSPT1 plasmid combined with 1 μg of pCAGGS-VP35-HA or pCEZ-VP30 (per well) were added to the co-transfection mix using the Lipofectamine 3000 transfection reagent. Two days later, transfected cells were washed with DPBS and lysed in 1X RIPA buffer including protease inhibitor cocktail (cOmplete™, EDTA-free Protease Inhibitor Cocktail), on ice for 15 mins. Soluble whole cell lysates were clarified by centrifugation at 12,000X rpm for 10 mins. FLAG-M2 affinity beads (40 μl per reaction) were washed with DPBS for once, NT2 buffer for twice before blocking with BSA in NT2 buffer (0.5 mg/ml) for 0.5 - 1 hour at room temperature. Lysates containing equal amounts of total protein (1-1.5 mg determined by BCA assay) were incubated with BSA-blocked FLAG-M2 affinity beads and rotated overnight at 4 ºC. Supernatant containing unbound proteins was removed by spin at 5000x g for 30 s, followed by four washes in 1 ml NT2 buffer. Immunoprecipitated proteins were eluted in 4X-SDS-loading buffer containing reducing agent with 10 mins incubation at 95 ºC and analyzed by western blotting.

### Viral growth kinetics and quantification of viral proteins and RNAs

Huh7 cells (5 × 10^4^ per well) were seeded in 24-well plates and transfected with 10 nM siRNA (NSC or GSPT1 si8 or UPF1si1, experimentally validated). Forty-eight hours after siRNA transfection, Huh7 monolayers were transferred into BSL-4 facilities and infected with EBOV-eGFP at specified MOI (PFU/ml) for one hour. Infection of the cells was followed by removal of the inoculum and replenishing with 2% FBS containing fresh media. In drug-treatment experiment, DMSO or CC-90009 (at specified concentration) containing 2% FBS-media was added to Huh7 monolayers (with the same seeding density) after removal of the virus inoculum. For virus titration, aliquots of supernatant were harvested at indicated time points from infected monolayers and used to infect Vero-E6 cells for foci-quantification. At desired time points, infected Huh7 monolayers were either stained with Hoechst and imaged by Olympus IX73 epi-fluorescent microscope using a 10X objective or collected directly in Trizol or 4x Laemmli buffer for inactivation and downstream RNA/protein quantifications. Inactivation procedures described here follow the approved standard operating procedures. Equal volume of Laemmli lysates were loaded on SDS-PAGE gel for western blot analysis. Equal amount of total RNA in each sample was used for strand-specific RT-qPCR. Fluorescent image acquired using Blue/Green Excitation for the GFP signal and Ultraviolet Excitation for the Hoechst-stained nucleus. Images from the same field of view but different channels were merge using ImageJ/FIJI.

### Strand-specific RT-qPCR

Total RNA was extracted from each Trizol-inactivated sample (per well) using Direct-zol RNA Miniprep Plus according to the manufacturer’s protocol, eluted in 50 μl DEPC-treated water and quantified by nanodrop. 200 ng of total RNA from each sample was used as the template for reverse transcription (RT) using High-Capacity cDNA Reverse Transcription Kit according to the manufacturer’s protocol. Strand-specific RT primer binding to the EBOV trailer region were used to amplify vRNA/genome. Total mRNAs, including EBOV mRNAs, were amplified using an Oligo-dT primer given the presence of poly-A tail. Resulting cDNAs were 1:100 diluted in DEPC-treated water and used in the subsequent quantitative real-time PCR on a Bio-Rad CFX Real-Time System. SYBR select master mix was used to mix with diluted cDNA template and target specific qPCR primer pairs in a 20 μl reaction. Each reaction was performed in triplicated wells on qPCR plates. PCR condition was following the Standard Cycling Mode (Primer Tm ≥ 60 ºC) according to manufacturer’s protocol. A default dissociation curve was performed immediately after the real-time PCR run to obtain the Tm (melting temperature) of each target. All qPCR primers specifically target EBOV sequence were confirmed by detecting no signal in MOCK infected control sample. The Tm of samples using the same qPCR primer pairs were confirmed are the same. Amplification plots have baseline subtracted, and the relative quantification (ΔΔC_T_) method was used to analyze results. GAPDH was used as the house-keeping gene for data normalization. In Figure 6B, 6G, 6H and 7C, samples from NSC were used as control to calculate the relative fold-change of the target amplicon in either GSPT1-knockdown (GSPT1si8) or UPF1-knockdown (UPF1si1) samples. In Figure 6H and 7F, samples from MOCK infected cells were used as control to calculate the relative fold-change of the target amplicon in EBOV infected cells. Sequence information of each primer been used is curated in **Table S3**.

## QUANTIFICATION AND STATISTICAL ANALYSIS

Statistical significance was assigned with P values were < 0.05 (*, *p < 0*.*05*; **, *p <0*.*01*; ***, *p <0*.*001*; ****, *p <0*.*0001*). If there was no statistical significance for a given comparison, it was not indicated in Figure 6F and 6G, but was indicated elsewhere as “ns”. Statistical details of experiments, including the type of test used, values and error bars displayed, the number of independent experiments, and the number of samples/technical replicates, can be found in all figure legends. Except analysis performed in proximity proteomics was described in “Bioinformatic analysis of mass spectrometry data” under “Method details”.

In summary, statistical analyses used in bar graphs comparing different conditions with the same control were either Welch’s one-way ANOVA or ordinary one-way ANOVA, both with Dunnett’s multiple comparisons test. For bar graphs with comparisons between two conditions (i.e., NSC to KD, with virus to without virus), Welch’s t test was used to determine the statistical significance of differences. Normality tests were performed to determine if the data are normally distributed. For all viral growth kinetics experiments, we first log-transformed viral titers prior to statistical analysis because the transformed data but not the raw data followed a normal distribution. We performed two-way, repeated measure ANOVA or mix-effect analysis (due to missing values) on the log-transformed viral titer data. Missing values were from those viral titers measured at one day post infection, which were below detection limit of the titration assay. Statistical significances describing differences as a matter of time and treatment were labeled at the end of all data lines, those describing differences of the treatment were labeled at the legend, and those describing the multiple comparisons at each time point were labeled on top of each data point. GraphPad Prism 9 was used.

## Notes

### Competing Interest Statement

The authors have declared no competing interest.

