## Supplemental Information for "Functional interactomes of the Ebola virus polymerase identified by proximity proteomics in the context of viral replication"

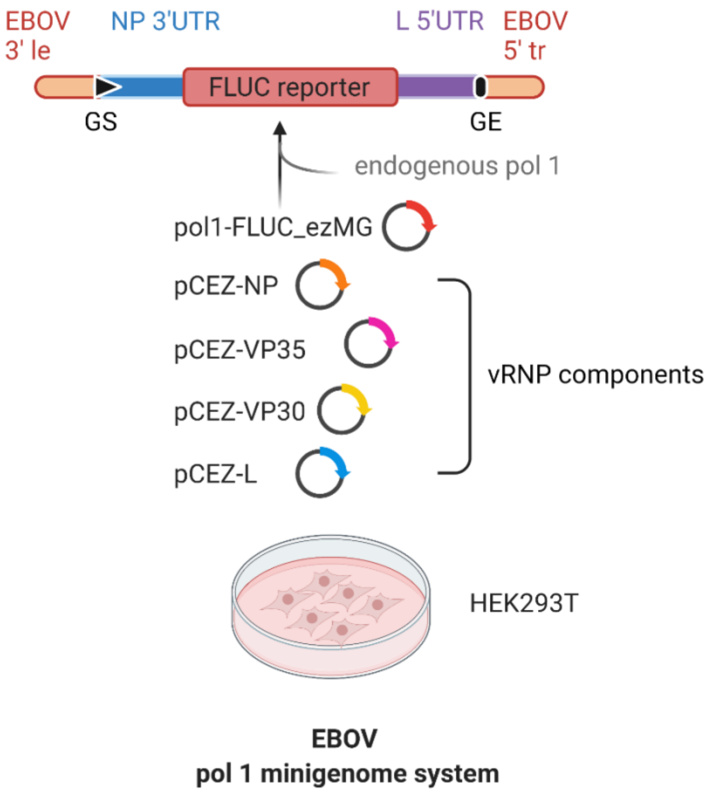


**Figure S1. Schematics of EBOV minigenome systems. Related to Figure 1.** Transfection of a plasmid encoding Firefly luciferase (Fluc) reporter flanked by the 3′ and 5′ extra-cistronic regions of authentic EBOV genome can produce a negative-sense, reporter viral RNA via endogenous polymerase 1-dependent transcription. Once transcribed, the reporter minigenome (ezMG) can be recognized, replicated or transcribed by co-expressed EBOV polymerase (L-VP35 complex) in the presence of EBOV NP and VP30 proteins. 3'le: Ebola Zaire leader sequence. 5'tr: Ebola Zaire trailer sequence. GS: gene-start signal of NP. GE: gene-end signal of L. UTR: untranslated region.

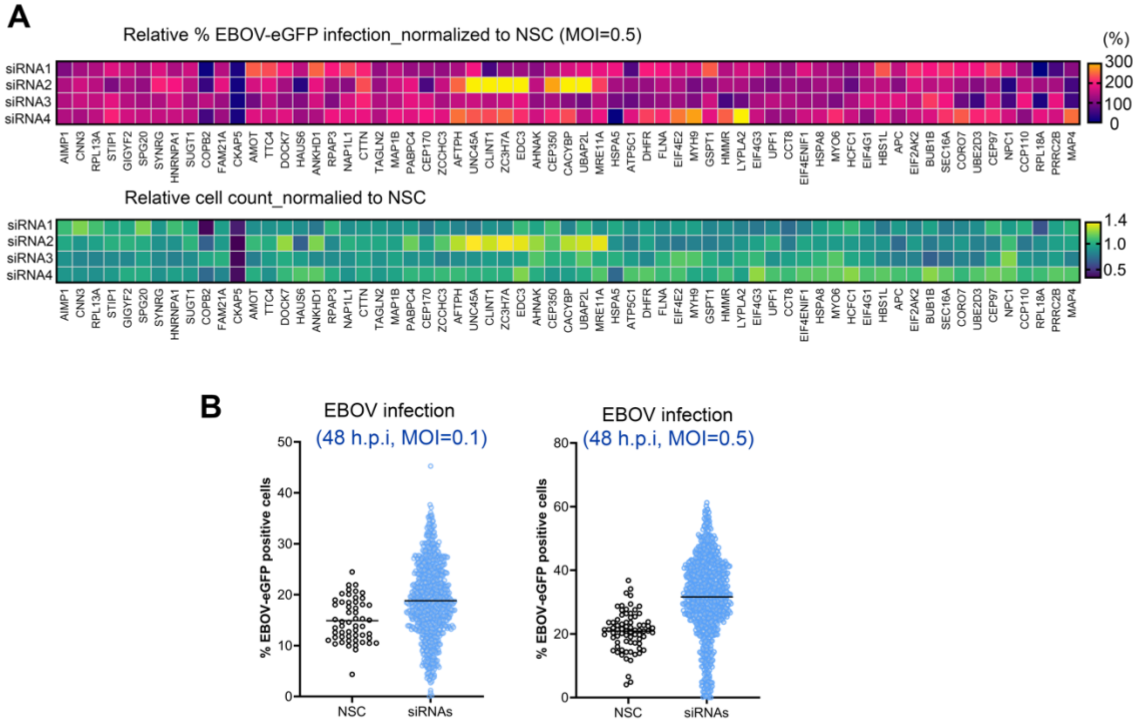


**Figure S2. EBOV infection and cell counts determined in the siRNA screens with raw infection rates of EBOV-eGFP in Huh7 cells. Related to Figure 3: repeated at a different MOI. (A)** Raw percentage of EBOV infection and cell count of each siRNA treatment were normalized to that of non-silencing controls, respectively. Normalized values are displayed in heat maps of relative percentage of infection and relative cell count. Each value is the mean of technical triplicates. Data points in %infection heat map with values exceeding the plotted range are marked in bright yellow. **(B)** The absolute percentage of cells infected with EBOV in the siRNA screen was quantified using CX5 high-content imaging system. To visualize the raw infection rate of each virus in Huh7 cells transfected with siRNA, the quantified % infection value corresponding to each well transfected with non-silencing control (NSC) or a given siRNA is pooled and plotted for each siRNA screen using a different MOI from Figure 3. MOI: Multiplicity of infection (PFU/cell); dpi: Day post infection.

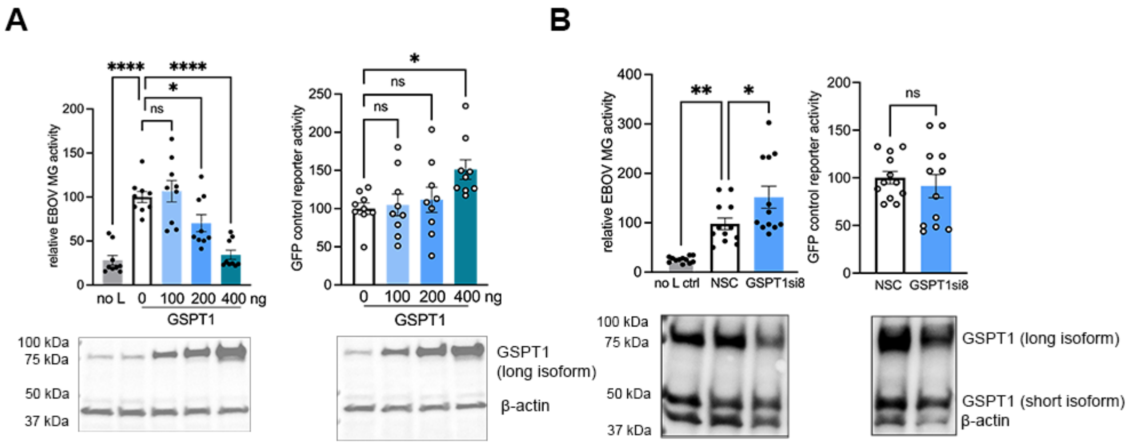


**Figure S3. GSPT1 is functionally relevant to EBOV RNA synthesis. Related to Figure 5.** Relative EBOV minigenome (MG) activity and a control GFP reporter activity in HEK 293T cells overexpressing GSPT1 **(A)** or transfected with a validated siRNA targeting GSPT1 (GSPT1si8) **(B)**. MG and reporter activity of the GSPT1-overexpressing/depleted cells are normalized to that of control cells. Raw luciferase activities and GFP intensities were first normalized to the corresponding amount of total protein in each sample to account for loading bias. Results from three (A) or four (B) independent experiments with triplicates were analyzed by using one-way ANOVA or Welch's t test under each condition compared to controls (ns, not significant; *, $p < 0.05$; **, $p < 0.01$; ***, $p < 0.001$; **** $p < 0.0001$). Values are means ± SEM. Representative western blot showing the expression levels of GSPT1 in different experimental conditions. Lysates from triplicate wells with the same experimental conditions were pooled (by equal volume) and analyzed.

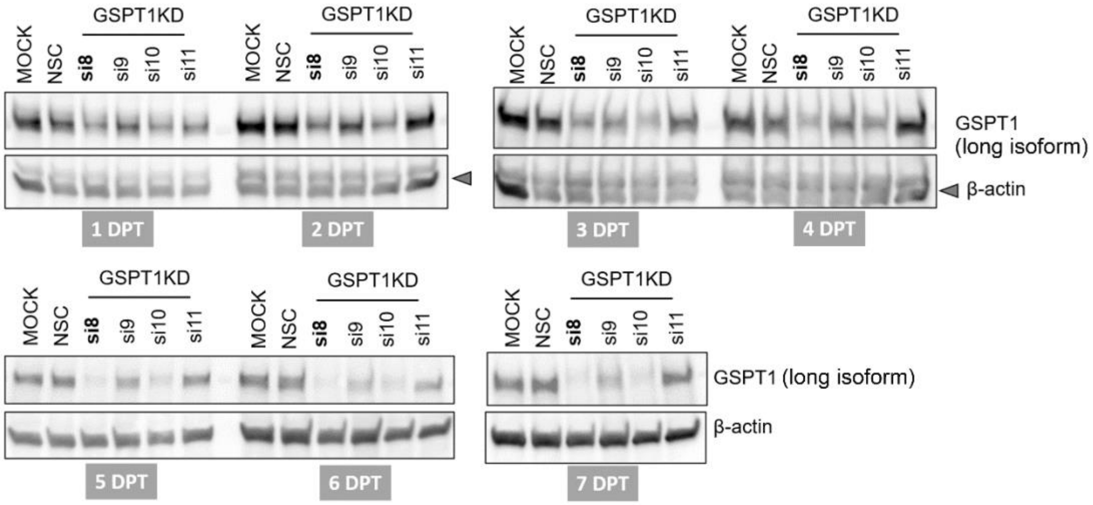


**Figure S4. GSPT1 knockdown kinetics mediated by siRNAs in Huh7 cells. Related to Figure 6.** Levels of endogenous GSPT1 protein (long isoform) determined by western blot in Huh7 cells upon individual siRNA-knockdown from one to seven days post transfection (DPT). Controls included non-transfected cells and cells transfected with the non-silencing control (NSC) siRNA. Four individual siRNA (si8 -10) were evaluated independently and si8 was selected for GSPT1 knockdown. For each lane, whole-cell lysates from triplicated wells were pooled and analyzed.

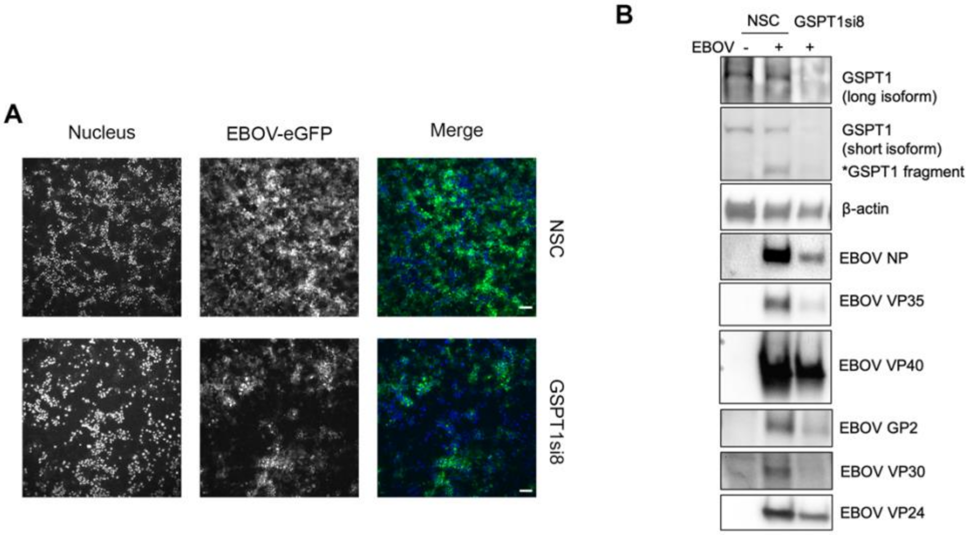


**Figure S5. EBOV-infected Huh7 monolayers and EBOV protein accumulation in GSPT1-depleted Huh7 cells. Related to Figure 6. (A)** A different biological replicate, EBOV growth curve experiment with GSPT1 KD, representative images of EBOV-infected Huh7 monolayer at 4 dpi are shown. Scale bar: 100 μm. **(B)** From the repeated experiment, western blot analysis of GSPT1 protein, loading control (β-actin), and viral protein levels in lysates from EBOV-infected Huh7 cells harvested at 4 dpi. Equal volumes of inactivated lysate were pooled from triplicates with the same treatment. Due to the low expression level of L protein, we could not consistently detect full-length L in inactivated lysate.

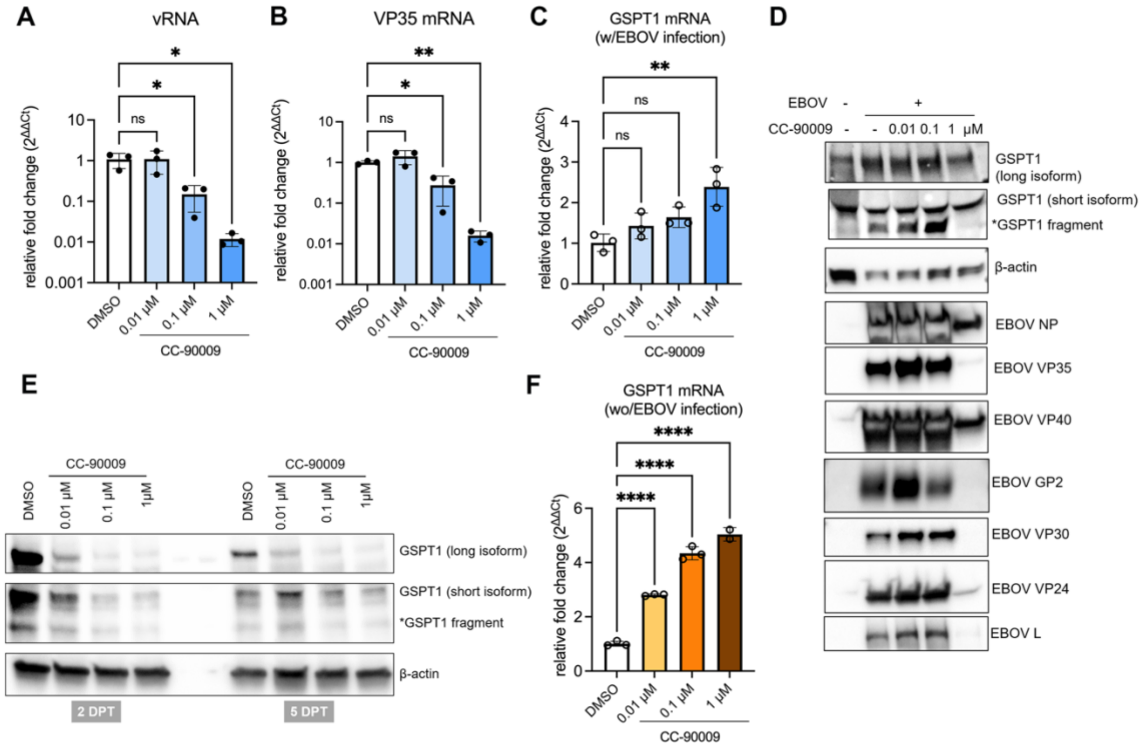


**Figure S6. Pharmacological targeting of GSPT1 protein strongly inhibits EBOV infection. Related to Figure 6.** Effect of CC-90009 treatment on **(A)** EBOV vRNA accumulation, or **(B)** mRNA accumulation at 4 days post-infection (dpi). Level of VP35 mRNA was used as a representation of EBOV mRNA level. Other EBOV mRNAs were similarly affected by CC-90009 treatment. **(C)** Levels of endogenous GSPT1 transcripts upon CC-90009 treatment in EBOV-infected Huh7 cells, at 4 days post-infection (dpi). **(D)** Western blot analysis of GSPT1 protein, loading control (β-actin), and viral protein levels in lysates from EBOV-infected Huh7 cells harvested at 5 dpi. Equal volumes of inactivated lysate were pooled from triplicates with the same treatment. **(E)** Levels of endogenous GSPT1 proteins in Huh7 cells upon CC-90009 treatment in the absence of viral infection. DMSO treatment serves as a control. Huh7 lysates were collected at 2- or 5- day post treatment (DPT) for western-blot analysis. **(F)** Levels of endogenous GSPT1 transcripts upon CC-90009 treatment in Huh7 cells, at 4 days post-treatment. For all bar graphs, results from one experiment with triplicate wells were analyzed by ordinary one-way ANOVA with Dunnett's multiple comparisons test to determine the statistical significance of the difference between DMSO and CC-90009 treatment at different concentrations (ns, not significant; *, $p < 0.05$; **, $p < 0.01$; ***, $p < 0.001$; **** $p < 0.0001$). Each data point represents the mean of technical triplicates performed in qPCR using the same cDNA. Values are means ± SD.

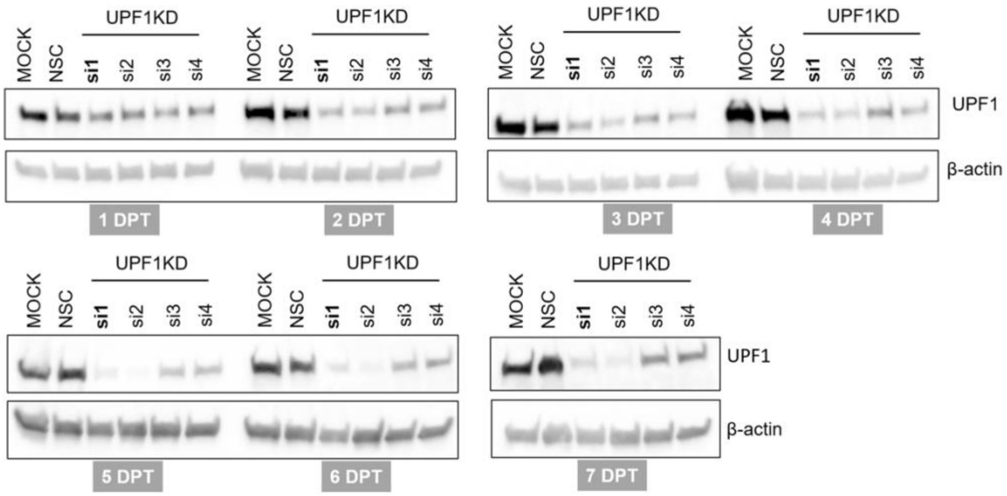


**Figure S7. UPF1 knockdown kinetics mediated by siRNAs in Huh7 cells. Related to Figure 7.** Levels of endogenous UPF1 protein determined by western blot in Huh7 cells upon individual siRNA-knockdown from one to seven days post transfection (DPT). Controls included non-transfected cells and cells transfected with the non-silencing control (NSC) siRNA. Four individual siRNAs (si1 - 4) were evaluated independently, and si1 was selected for UPF1 knockdown. For each lane, whole cell lysate from triplicated wells was pooled and analyzed.

**Table S1. Proteomic results. Related to Figure 2.** (Provided as a separate Excel file)

Relative abundance ratio and the corresponding P-value for each protein identified in all proximity-proteomic samples.

**Table S2. siRNA sequences. Related to Figure 3, Figure 5-7 and Figure S2-7.**

(Provided as a separate Excel file)

The sequence information of each individual siRNA used in the siRNA screen and used in GSPT1 or UPF1 knockdown.

**Table S3. Primers used in this work. Related to Figure 6, Figure 7, and Figure S6.**

| Primer name | Sequence (5' to 3') | Used for | Target |
|---|---|---|---|
| RT-EBOV-vRNA | CTCATAAATTGCTCTCATACATCATATTGA | Reverse transcription | EBOV vRNA (Trailer) |
| Oligo dT (20mer) | TTTTTTTTTTTTTTTTTTTT | Reverse transcription | Polyadenylated mRNAs |
| EBOV_vRNA_F | TACTGCCGCAATGAATTTAACG | qPCR | EBOV vRNA Trailer sequence |
| EBOV_vRNA_R | ATAACAATATGAGCCCAGACCTT | qPCR | EBOV vRNA Trailer sequence |
| EBOV_NP_F | CAGTGCGCCACTCACGGACA | qPCR | EBOV NP coding sequence |
| EBOV_NP_R | TGGTGTCAGCATGCGAGGGC | qPCR | EBOV NP coding sequence |
| EBOV_VP35_F | CTGGATCTCTGAGCAGCTAATG | qPCR | EBOV VP35 coding sequence |
| EBOV_VP35_R | CATTTGGGATGCGTAGCATAATC | qPCR | EBOV VP35 coding sequence |
| EBOV_VP40_F | GGCGGGTTATATTGCCTACT | qPCR | EBOV VP40 coding sequence |
| EBOV_VP40_R | GGTGTCAGGAAGCCTGTATT | qPCR | EBOV VP40 coding sequence |
| EBOV_VP30_F | CACTGATCAAGACGGCAGAA | qPCR | EBOV VP30 coding sequence |
| EBOV_VP30_R | CTCGTCATCACAGCACATAGAG | qPCR | EBOV VP30 coding sequence |
| EBOV_VP24_F | GCGACCTCTGTAACTTCTTAGT | qPCR | EBOV VP24 coding sequence |
| EBOV_VP24_R | TAGGGCCATTCCTTTGTGAG | qPCR | EBOV VP24 coding sequence |
| EBOV_L_F | CGCAACTACGCAACTGTAAAC | qPCR | EBOV L coding sequence |
| EBOV_L_R | CCTTGAGAAGAACTGGGACTATG | qPCR | EBOV L coding sequence |
| EBOV_VP35/VP40_F | TCATATCTCGCTAGCAGCTTAAA | qPCR | EBOV VP35 3'UTR |

| | | | |
|---|---|---|---|
| EBOV_VP35/VP40_R | TCTCAGCCGAGGTAGGTTT | qPCR | EBOV VP40 5'UTR |
| EBOV_VP40/GP_F | GGTTGTTCACAATCCAAGTACAG | qPCR | EBOV VP40 3'UTR |
| EBOV_VP40/GP_R | AGAGAGATGAAGATTACGCTCAC | qPCR | EBOV GP/sGP coding sequence |
| EBOV_VP30/24_F | GTGGAGGAGGTGTTTGGTATT | qPCR | EBOV VP30/VP24 intergenic region (IGR) |
| EBOV_VP30/24_R | TAGCATTTCCGGTCACAATCT | qPCR | EBOV VP30/VP24 intergenic region (IGR) |
| EBOV_VP24/L_F | CAGGTCACATGTGTTAGGTTTC | qPCR | EBOV VP24 3'UTR |
| EBOV_VP24/L_R | AAACACGGAAAGACCCAATAAG | qPCR | EBOV L 5'UTR |
| GSPT1_F | GAGGAGGAAGAGGAAATCCC | qPCR | All GSPT1 transcript variants |
| GSPT1_R | TCCTTTTGTCAACCATTCCA | qPCR | All GSPT1 transcript variants |
| UPF1_F | ACCTATTACACGAAGGACCTCC | qPCR | All UPF1 transcript variants |
| UPF1_R | ACGTCCGTTGCAGAACCAC | qPCR | All UPF1 transcript variants |
| GAPDH_F | CATGAGAAGTATGACAACAGCC | qPCR | GAPDH |
| GAPDH_R | TGAGTCCTTCCACGATACC | qPCR | GAPDH |
